# Development of an *Agrobacterium*-delivered codon-optimized CRISPR/Cas9 system for chickpea genome editing

**DOI:** 10.1101/2022.09.20.508560

**Authors:** Santosh Kumar Gupta, Niraj Kumar Vishwakarma, Paheli Malakar, Poonam Vanaspati, Nilesh Kumar Sharma, Debasis Chattopadhyay

## Abstract

Chickpea is considered recalcitrant to *in vitro* tissue culture. The Clustered, Regularly Interspaced Short Palindromic Repeats/CRISPR-associated protein 9 (CRISPR/Cas9) based genome editing in chickpea can remove the bottleneck of limited genetic variation in this cash crop rich in nutrients and protein. However, the generation of stable mutant lines using CRISPR/Cas9 requires efficient and highly reproducible transformation approaches. We modified a binary vector pPZP200 by introducing a codon-optimized Cas9 gene for chickpea and the promoters of *Medicago truncatula* U6 snRNA for expressing guide RNA targeted to the Phytoene Desaturase (PDS) gene. The dissected single cotyledons with half embryo of chickpea were used as explants for genetic transformation. A single gRNA was found sufficient to achieve high efficiency (42%) editing with the generation of PDS mutants with albino phenotypes. A simple, rapid, highly reproducible, stable transformation and CRISPR/Cas9-based genome editing system for chickpea was established. For the first time, this study aimed to demonstrate this system’s applicability by performing a gene knockout of the chickpea phytoene desaturase gene (*CaPDS*) in stable shoots using an improved chickpea transformation protocol.

## 1. Introduction

Chickpea (*Cicer arietinum* L.) is one of the grain legume crops domesticated in the old world (earlier than 9500 BC) and widely cultivated in more than 50 countries (Upadhyaya *et al.*, 2011). This staple legume crop is highly enriched in nutrients, fibre, and protein (Chibbar *et al.*, 2010). Conventional breeding has contributed to chickpea improvement, but many problems still need to be addressed. Genome editing might be a new approach that offers the maximum possible gateway for genetic modifications. The Clustered Regularly Interspaced Short Palindromic Repeats/ CRISPR associated protein 9 (CRISPR/Cas9) based genome editing in chickpea offers a promising tool for its genetic improvement such as abiotic stress (increased suppleness to drought, salt, heat, and cold tolerance), increased seed nutrient density, seed yield, biofortification of micronutrients and resistance to various biotic stress (nematodes, viruses, and fungal diseases). Generating stable mutant lines using CRISPR/Cas9 requires an efficient and highly reproducible transformation method. Tissue culture-based regeneration of chickpea is considered challenging (Bhowmic *et al.*, 2019; Varshney *et al.*, 2009). *Agrobacterium*-mediated chickpea transformation and regeneration is highly difficult among all leguminous crops. Advancing chickpea transformation can remove the hurdles of its limited genetic variation and bottlenecks for implementing high-throughput genome editing in this nutrient and vegetarian protein-rich cash crop.

Several reports on chickpea transformation are available, but none of these methods has high efficiency and virtuous regeneration capacity. Due to minimal *in vitro* rooting and hard-hitting hardening steps of transformed positive tissue cultured plants, very low transformation efficiency, poor success rate, and non-transmission of genes to subsequent generations were recorded (Popelka et al., 2004; Krishnamurthy et al., 2000; Sarmah et al., 2004; Sanyal et al., 2005; Bhowmik et al., 2019). We have developed an advanced, modified, simple, and efficient protocol for chickpea transformation, regeneration, selection, and hardening of the micro-grafted and rooted plants. This protocol is robust and highly reproducible. We have side-stepped many earlier reported steps during protocol optimization studies. The shortening of optimization steps helps reduce the tissue culture period and avoid somaclonal variation and contamination in transgenic plants. We have also optimized the micrografting steps and found that micro-grafting of *in vitro* regenerated shoots onto homozygous rootstock grown *in vivo* proved to be an uncomplicated and consistent method for establishing transgenic plants. A strong natural rooting system development has been optimized in transformed shoots devoid of growth regulators and selection markers. We have reproduced our optimized chickpea transformation and regeneration protocols with different constructs.

CRISPR/Cas9-based gene-editing methods have been successfully established for various legumes like soybean (Liu *et al.*, 2019), cowpea (Ji *et al.*, 2019), and model legumes such as *Medicago truncatula* (Meng *et al.*, 2017) and *Lotus japonicus* (Wang *et al.*, 2019). However, the recalcitrance nature of *in vitro* gene transfer and regeneration has posed a serious challenge to gene editing and the generation of stable knockout plants in chickpea (Bhowmic *et al.*, 2019). A single report is available based on cell-free protoplast transformation utilizing CRISPR/Cas9 DNA-free gene editing of a drought tolerance-associated gene in chickpea at a cellular level (Badhan et al., 2021). However, theoretical approaches lack this versatile tool’s experimental demonstration and applicability to generate full-grown knockout plants. Therefore, a scorable marker gene is required to validate any new strategy for generating CRISPR/Cas9-based knockout plants. In tomatoes, the Phytoene desaturase gene (*PDS)* was used as a common marker gene for the detection of editing using VIGS (Virus-induced gene silencing) (Liu *et al.*, 2002). Knockout of the *PDS* gene by CRISPR/Cas9 moderated chlorophyll, carotenoid, and gibberellin biosynthesis leading to albino and dwarf phenotype (Qin *et al.*, 2007). This study has been reported in melon (Hooghvorst *et al.*, 2019), Medicago, cucumber (Chandrasekaran *et al.*, 2016), watermelon (Tian *et al.*, 2016), and hybrid poplar (Bae *et al.*, 2021) with a single guide RNA (gRNA). This scorable marker, like the PDS gene, can better validate the feasibility of a gene-editing method as its disruption causes photobleaching or the albino phenotype in the transformed plants. We targeted the chickpea *Phytoene Desaturase* gene (*CaPDS)* for demonstration of the capability of our construct and the method. A chickpea codon-optimized *Streptococcus pyogenes* Cas9 (*spCas9)* gene and newly assembled pPZP200 binary vector were used in this study (_pro_CaMV35::Cas9). The Albino phenotypes with high-efficiency editing were achieved in transformed, and tissue culture regenerated plants. A single gRNA was found sufficient to generate the knockout plants in chickpea.

## 2 Materials and methods

### 2.1 Plant materials and culture media

To establish an improved protocol for chickpea transformation, mature seeds of chickpea desi cultivar Pusa-362, BGD-72, ICC4958, DCP92-3, ICCX-810800, and Kabuli ICCV2 were obtained from the Indian Agricultural Research Institute (IARI) New Delhi, Indian Institute of Pulses Research (IIPR) Kanpur, and ICRISAT, Hyderabad and multiplied at the experimental fields of NIPGR, New Delhi and SVPUAT Meerut. The media composition used in this study contained MS basal medium (Murashige and Skoog, 1962) supplied by Duchefa Biochemie, including B5 vitamins (Gamborg et al., 1968) by Sigma. The pH of all tissue culture media used in this study was 5.8 before adding agar and proceeding to the autoclave. All selective agents were sterilized using a 0.45μm filter. 3% sucrose (Sigma) and 0.8% (w/v) agar (Duchefa Biochemie) were used to supplement all the tissue culture media except co-cultivation media, devoid of sucrose and agar.

### 2.2 *Agrobacterium* tumefaciens strains

*Agrobacterium tumefaciens* strains GV3101, LBA4404, and EHA105 were used in this study. We found that GV3101 exhibited better efficiency than the other two strains. For further chickpea transformation studies, GV3101 was used. *Agrobacterium* strains harbouring the binary vector were streaked using glycerol stock onto LB agar plates containing Rifampicin (50 mg L^-1^) and kanamycin (50 mg L^-1^) and incubated for two days at 28 °C.

### 2.3 Cloning of sgRNA into a vector

The single guide RNA target was designed using CHOPCHOP sgRNA design online tool (http://chopchop.rc.fas.harverd.edu/) (Montague et al., 2014). CHOPCHOP predicts the frameshift rate of all the targets for high-efficiency editing with the predicated score of possible off-target. In this study, a 20-nucleotide single guide RNA of CaPDS gene was cloned under the MtU6.1 promoter with a SpCas9-specific conserved scaffold sequence of 76 nucleotides and a MtU6.1 terminator sequence. The guide RNA and scaffold-infused fragments (sgRNA-scaffold) were amplified from a custom synthesized cassette (Thermo Fisher Scientific, Inc.). Four sets of primers were used to synthesize a double-stranded sgRNA expression cassette with an overlapping strategy for different targets (Supplementary Table 1). The construct is flanked by suitable restriction sites (Pac1 & Asc1) in their respective forward and reverse primers. These restriction sites were further used to assemble the sgRNA1 cassette into the Cas9 binary vector.

### 2.4 *A. tumefaciens-mediated* transformation and regeneration; improved protocol

#### 2.4.1 Seed sterilizations ad Explant Preparations

Seeds were washed 2-3 times with distilled water and kept with Tween-20 (1%) for 10 min under stirring conditions. Then, these seeds were further rinsed with autoclaved distilled water 2-3 times and sterilized with 1% sodium hypochlorite solution for 15 min. After this, the seeds were washed with autoclaved water (3-4 times), 100% ethanol (2-3 times), autoclaved water (3-4 times) consecutively, and finally soaked for 8-10 hours (h) for desi cultivar and 5-6 h for Kabuli cultivars in the dark.

#### 2.4.2 Explant Preparation

After removing water from the overnight soaked seeds, the seed coat was detached; the radicle was removed. Then the seeds were cut longitudinally along the plumule with the help of a scalpel under sterilized conditions. The separated half cotyledons with half embryo chickpea seeds were selected as explants for transformation. These explants were further utilized for *Agrobacterium* tumefaciens infection. Explants were prepared from chickpea seeds in different independent transformation events for half cotyledon and half embryo type explants. From overnight soaked chickpea seeds of cultivar ICC4958, mature embryo explants were prepared and transformed.

#### 2.4.3 Plant transformation and Regeneration events

The explants obtained were pre-incubated aseptically for 24 h on MS basal medium (with vitamins and sucrose). A single colony was used in a culture in an LB media containing Rifampicin (50 mg L^-1^) and Kanamycin (50 mg L^-1^) and incubated at 28°C at 180 rpm. This primary culture was used to inoculate (0.1% of overnight grown culture) secondary culture into 50 mL of LB media containing Rifampicin (50 mg L^-1^) and Kanamycin (50 mg L^-1^). 50 μL of 100 mM acetosyringone was added after the OD (optical density) of the secondary culture reached 0.6 to 0.8. After centrifugation at 5,000 rpm for 5 min, the culture was resuspended in a resuspension medium containing 100 μM acetosyringone. Again, after centrifugation, the supernatant was discarded, and the precipitated cells were washed with 15 ml of sterile half-strength MS followed by centrifugation at 5,000 rpm. The precipitated cells were resuspended into a 30 ml half-strength MS basal liquid medium for further use in co-cultivation. Explants were dipped for exposure of *Agrobacterium* strain GV3101 harbouring plant transformation vectors pBI121, pGWB2-modified (OD_600_=0.8) for 30 minutes (Supplementary Text 2). After removing an excess bacterium by blotting on sterile paper, explants were co-cultivated for 48 h in a co-cultivation medium (MS basal medium supplemented with vitamins and 3% sucrose) at 22°C in the dark.

Following co-cultivation, the explants were transferred to Shoot Regeneration Medium (SRM-1) fortified with Cefotaxime (250 mg L^-1^) and the first selection of Kanamycin (50 mg L^-1^) for 7-10 days under continuous cool and white light fluorescent lamps devoid of growth regulators. The regenerated small shoots were further shifted to SRM-2, which was invigorated with Cefotaxime (250 mg L^-1^) and the second selection of Kanamycin (100 mg L^-1^) for 12-15 days. After 15 days, healthy shoots have appeared in most of the explants. These explants were further shifted on the third set of selection fortified with Cefotaxime (250 mg L^-1^) and Kanamycin (200 mg L^-1^) SSM for the next 12-15 days. During this period, the shoot induction was completed. During the sub-culturing of the emerging shoots, untransformed and bleached yellowish light green shoots were removed. Then, the healthy green shoots were selected for micro-grafting. An average of 60-65% of strengthened shoots survived well as healthy plants. A cluster of well-developed shoots was separated and further shifted onto Shoot Maintenance Medium (SMM) (Supplementary Figure 2). These shoots were further used for micro-grafting and the establishment of mature transgenic plants. This selection step was repeated 2-3 times for 7-10 days.

### 2.5 Detection of transgene and CRISPR/Cas9 mutation

Leaflets from transformed T0 and wild-type (non-transformed) control plants were collected, and genomic DNA was isolated using the CTAB method as previously described (Saghai-Maroof *et al.*, 1984). Each T0 plant was amplified with specific primers for NPTII and Cas9 to confirm the presence of transformed plants by the designed construct. In addition, a fragment flanking gRNA (256bp) of the *CaPDS* gene from transgenic chickpea plants was amplified using suitable primers (Supplementary Table 1), and Sanger-sequencing gel-purified the amplified PCR products were used for the detection of specific additions, deletions, and substitutions. Finally, the mutation rate was calculated based on mutated and non-mutated transformed samples collected from albino and green plants.

## 3 Results

### 3.1 Development of an improved genetic transformation method for chickpea

Earlier reported chickpea transformation methods were associated with low transformation efficiency, poor in vitro rooting, hardening, and establishment of mature plants (Huda et al., 2000; Sarmah et al., 2004; Polowick et al., 2004; Senthil et al., 2004; Sanyal et al., 2005; Chakraborti et al., 2006; Bhatnagar-Mathur et al., 2009; Bhowmik et al., 2019). To overcome these hurdles in chickpea transformation, we have used a modified protocol for chickpea transformation. Single cotyledon with half embryo explants were prepared from overnight soaked chickpea seeds. These explants were exposed to three different recombinant *Agrobacterium* strains GV3101, EHA105 and LBA4404 for transformation (Table 2; Supplementary Tables 2 & 3). During transformation optimization studies, GV3101 showed better efficiency than the other two strains. The media composition was modified (devoid of growth regulators) for the regeneration of shoots, which are designated as shoot regeneration media (SRM-1 & SRM-2). The explants and shoots were incubated at 22±2°C under cool white light with an intensity of 60 μmole m^-2^ s^-1^ under a time regime of 10 h light and 14 h dark. The sub-culturing was performed at regular intervals (3 cycles of 07-10 days). The composition of the medium was optimized to reduce the time, labour, and cost of tissue culture in chickpea transformation (Table 1). The binary vectors pBI101.2 and modified pGWB2 harboring β-glucuronidase (GUS) and Green fluorescent protein (GFP) under the *CaMV35S* promoter were used in the transformation, respectively. Transformed and regenerated plants were screened by endpoint PCR using GUS and GFP-specific primers (Supplementary Figure 1). Different parts (Cotyledon and embryo, seed coat, pod cover, flower, root, and leaves) of the regenerated plants were exposed for histochemical detection of GUS assay. Transformed and regenerated chickpea plants carrying a Nuclear Localization Signal-Green fluorescent protein (NLS-GFP) gene under *CaMV35S* promoter were also developed. GFP was observed in the nucleus by confocal microscopy (Leica SP8, Leica Microsystems) in the leaves of transformed plants (Figure 1A).

**Figure 1:**
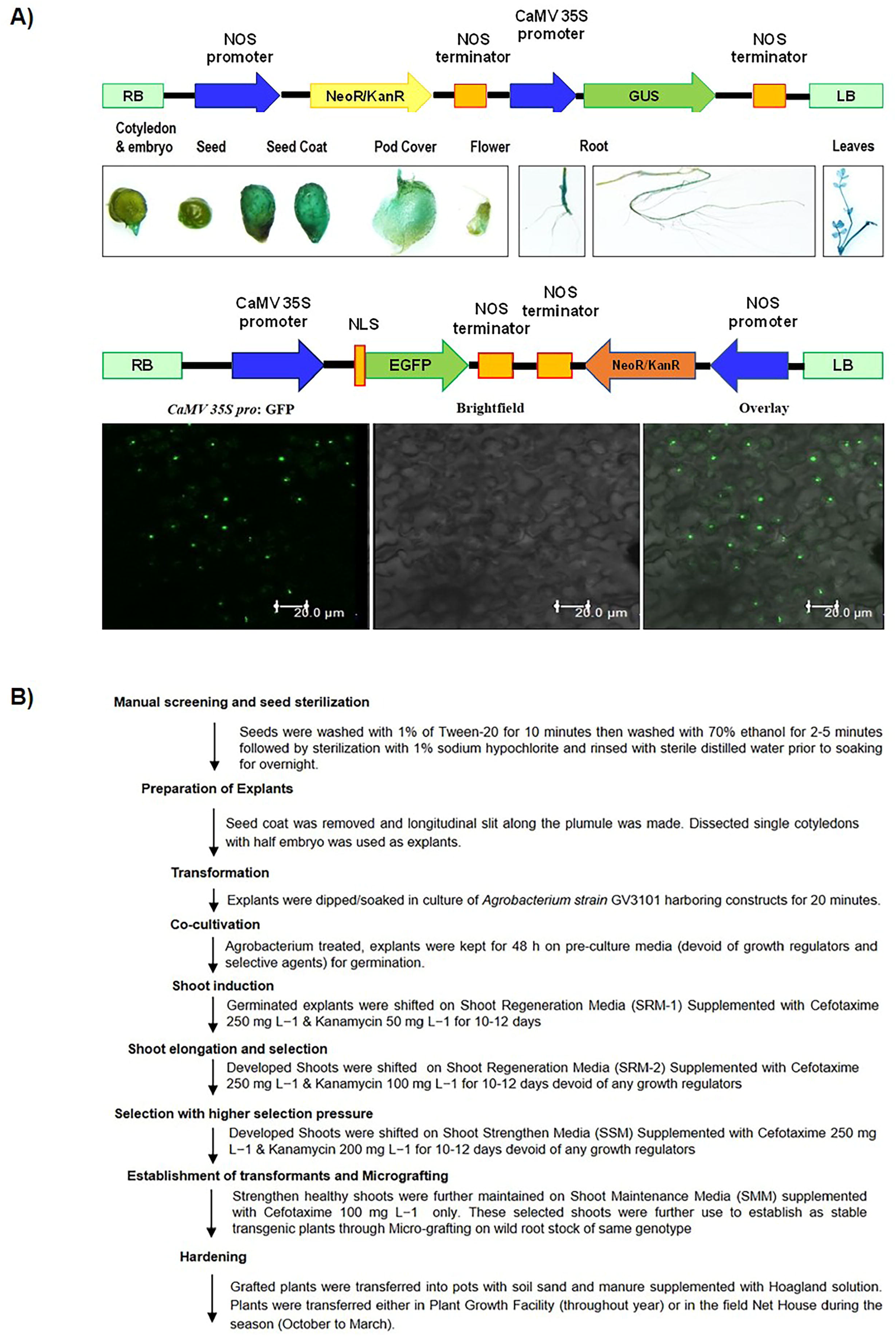
Plant transformation vectors and an improved protocol for *Agrobacterium*-mediated step-wise schematic description of chickpea transformation. **(A)** Plant transformation vectors containing β-glucuronidase gene (*GUS*) (pBI101) and Green fluorescent protein (*GFP*) gene (pGWB2-modified) driven by CaMV 35S promoter and NOS terminator were used for the development of transgenic plants under the selection marker Kanamycin. Hardened and established PCR-positive GUS lines were evaluated for the tissue-specific expression in vegetative (root and leaves) and reproductive parts (cotyledon and embryo, seed, seed coat, pod cover, and flower) chickpea plants. GFP-derived fluorescence, bright field, and merged image detected by laser scanning confocal microscope (TCS SP8, Leica, Microsystems, Germany) in transformed plant leaves. The scale bar represents 20μm. **(B)** Schematic representation of the steps required for *Agrobacterium tumifaciens*-mediated chickpea transformation.

**Table 1:** Details of media used in Chickpea transformation and regeneration.

| S N. | Media | Components | pH | Remarks |
| --- | --- | --- | --- | --- |
| 1. | SRM-1 | MS 4.4 gL<sup>-1</sup>, Cefotaxime 250mgL<sup>-1</sup>, kanamycin 50 mgL<sup>-1</sup>, 0.8% Agar | 5.8 | 10 days storage |
| 2. | SRM-2 | MS 4.4 gL<sup>-1</sup>, Cefotaxime 250mgL<sup>-1</sup>, kanamycin 100 mgL<sup>-1</sup>, 0.8% Agar | 5.8 | 10 days storage |
| 3 | SSM | MS 4.4 gL<sup>-1</sup>, Cefotaxime 250mgL<sup>-1</sup>, kanamycin 200mgL<sup>-1</sup>, 0.8% Agar | 5.8 | 10 days storage |
| 4 | SMM | MS 4.4 gL<sup>-1</sup>, Cefotaxime 250mgL<sup>-1</sup>, 0.8% Agar | 5.8 | 10 days storage |

**Table 2:** *Agrobacterium tumifaciens* (GV 3101) mediated the transformation efficiency of chickpea cv. ICC4958.

| Construct used | Explant type | No. of explant | No. of regenerated shoots (After cocultivation) | No. of a kanamycin-resistant single shoot on different concentration |  |  |  |  |  | Average Transformation n frequency (%) |
| --- | --- | --- | --- | --- | --- | --- | --- | --- | --- | --- |
|  |  |  |  | Kan 50 |  | Kan 100 |  | Kan 200 |  |  |
|  |  |  |  | I | II | I | II | I | II |  |
| ProCaMV::GUS | Half embryo longitudinally with one cotyledon | 1850(15) * | 1080 | 890 | 795 | 683 | 611 | 498 | 380 | 20.54 |
| ProCaMV::GFP | Half embryo longitudinally with one cotyledon | 1498(12) * | 820 | 780 | 680 | 501 | 308 | 248 | 208 | 18.09 |
|  | Total | 3348(27) * | 1900 | 1670 | 1475 | 1184 | 919 | 746 | 588 | 17.56 |

\*No. of independent transformation events

A total of 1850 single cotyledon and half embryo explants were prepared from 950 chickpea seeds in 15 independent transformation events for the *_Pro_CaMV::GUS* and 1498 explants for the construct *_Pro_CaMV::GFP* marker genes to check the efficiency of the modified transformation protocol in 12 independent events. During the preparation of explants and exposure to recombinant *Agrobacterium* culture, we have used the resuspension medium devoid of sucrose and half-strength MS basal with pH 5.5. The initial co-cultivation of explant and culture was optimized for the specific period and incubation conditions. For better interaction, a 30 minutes exposure under shaking (72 rpm) incubation condition at 28°C. These explants were dried on autoclaved blotting sheet to remove excess culture and medium.

Further, these explants were transferred on MS basal medium supplemented with vitamins and 3% sucrose. After 72 h, these explants were transferred to shoot regeneration media (SRM-1) for regeneration and selection. The growth media used for tissue regeneration and selection were devoid of growth regulators. Out of 1850 and 1498 single cotyledon and half embryo explants, 1080 and 820 had shoot buds in ProCaMV::GUS and ProCaMV::GFP, respectively, on SRM-1. This observation was recorded after seven days. At this stage, morphogenic changes of the cultured explants were recorded. The initial selection pressure of the selection marker was lowest (Kan 50mg L^-1^, SRM-1) and doubled in the next stage (Kan 100mg L^-1^, SRM-2) in two passages of 8-10 days intervals for both the medium, respectively. These two passages recorded 890 and 780 shoot buds containing explants in _Pro_CaMV::GUS and _Pro_CaMV::GFP, respectively. Further, in this method, subcultures of the explants on the Shoot strengthen medium (SSM) with the highest selection pressure (Kanamycin 200 mg L^-1^) result in multiple regenerated healthy chickpea shoots. Kanamycin-selected shoots were maintained onto Shoot Maintenance Medium (SMM) (Table 1). Shoots maintained on SMM were further used for micro-grafting and rooting to establish mature transgenic plants (Figure 1B; Supplementary Figure 2). The total regeneration frequencies for both the constructs were calculated as 20.54% and 18.09% in _Pro_CaMV::GUS and _Pro_CaMV::GFP, respectively (Table 2).

### 3.2 Micrografting of regenerated transformed shoots on untransformed rootstock

To establish the positive transformants of chickpea, a modified mid-grafting method was used. This method selected the *in vitro* derived transgenic shoots (scion) and soil-raised homozygous wild rootstock (6-12 days old) of the same genotype. To avoid any fungal infection, the soil was prepared in the ratio of 3:1:1 Soil-Sand-Vermiculite mixture. The diagonal cuts on scion and rootstock were made using a new and bench sterilized surgical blade. Approximately the girth of the plant was 0.8 cm in diameter. About 1-1.5 cm long chickpea plantlets were chosen. Creating a smooth and sloping cut in scion at one stroke is required. The cut was made in a reverse downwards direction in the rootstock, starting about ¼ of the distance from the top. The scion and rootstock were locked properly. Supporting material of silicone polymer ring (Sigma) or coating with petroleum jelly was used to stabilize this assembly. This assembly was slid to a combination point of scion and rootstock to provide appropriate strength. The scion was notched from both sides, removing the skin of the stock. The healthy rootstock was grown in a small pot. After completing micrografting steps (1-10), the pot was watered and properly covered with a polybag to maintain the humidity (Figure 2). A regeneration media (2-3 days old only), precise pH (5.6-5.8), and standard chickpea tissue culture plant growth condition (22±2°C under a time regime of 10 h light and 14 h dark, 60% relative humidity, with a light intensity of 150 μEm^-2^ s^-1^) was maintained. The plants were watered with ½ MS or ¼ Hoagland solutions per their requirement.

**Figure 2:**
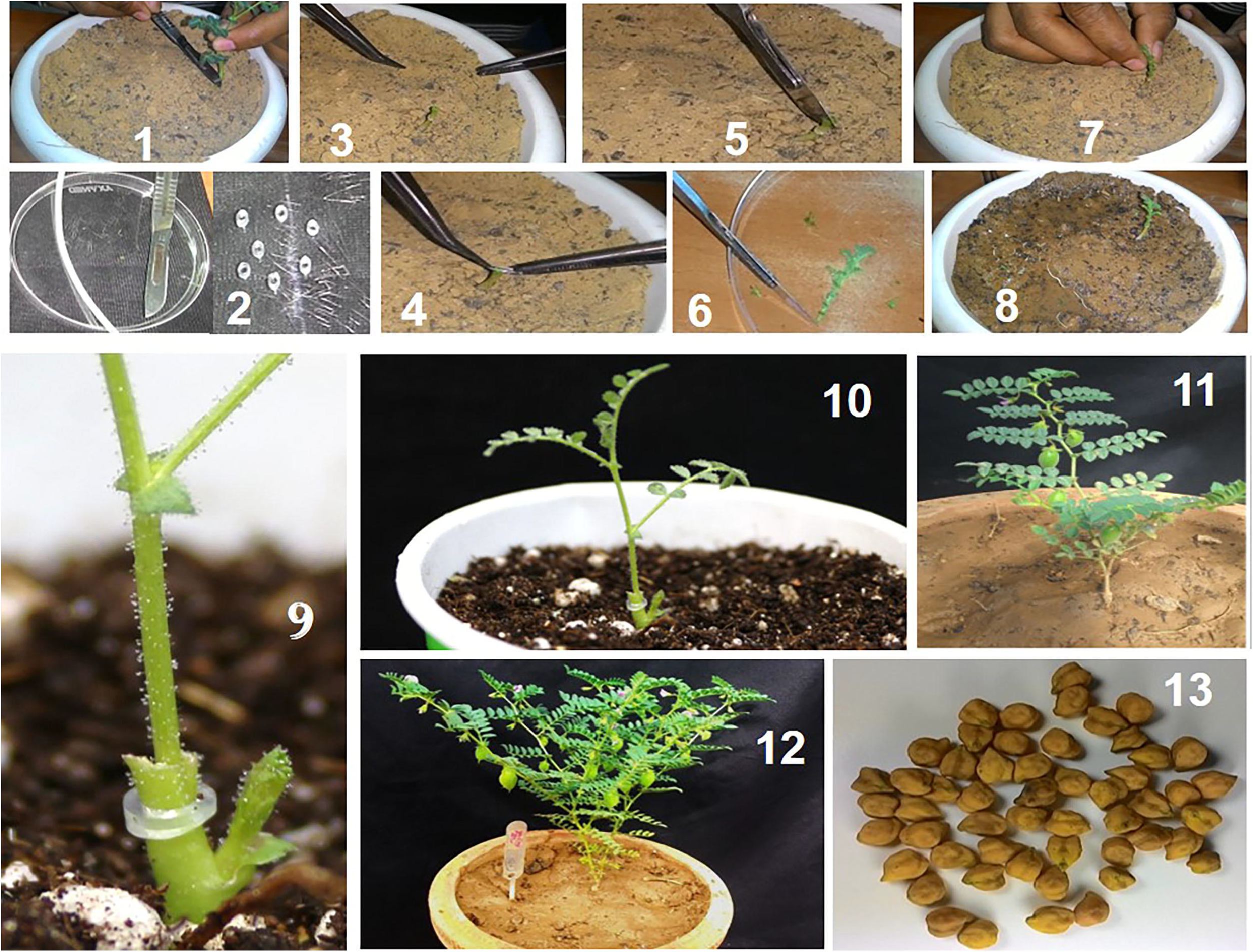
Establishment, hardening, and evaluation of transformed chickpea plants. **(A)** Establishment of regenerated and selected shoots through micro-grafting and hardening. Micro-grafting of putative transgenic shoots on wild rootstock through mid-grafting approach (1-10). Soil hardening of grafted plants. After 15 days, soilrite hardened plants were further shifted to garden soil: sand: manure mixture (3:1:1) is a comparatively larger pot for flowering and maturity (11-13).

### 3.3 Induction of natural roots from the transformed shoots

This alternative speedy procedure is based on generating natural roots from the transformed and regenerated shoots. The use of antibiotic selection is avoided in this method. After shoot regeneration on SRM-1 media, the regenerated shoots were directly shifted to the final selection rooting media (CRM; ½ MS 2.2 gL^-1^, Cefotaxime 250 mgL^-1^ and 0.8% Agar). Then, the rooted plants were transferred onto the soil medium (3:1:1, Agropeat, soilrite, and vermiculite)-filled paper cups. These plants were kept for four days in controlled conditions.

Further, these plants were transferred onto pots filled with specific garden soil, sand, and manure (3:1:1) in standard chickpea growth conditions, covered with appropriate polythene bags, and grown to maturity. The polythene bags are removed after the hardening of the plants. The presence of the transgene was tested in the next generation. A total of 291 regenerated healthy shoots were selected for in vitro rooting using CRM in 30 independent events. We have recorded 54.98% (survival of 160 positive plants out of 427 regenerated shoots) survival of the putative transformants as a plant by this method (Supplementary Figure 3).

### 3.4 Target selection and sgRNA design and expression cassettes

We have chosen to disrupt the PDS gene of chickpea (*CaPDS; 15-cis-Phytoene Desaturase)* located in chromosome 5, having a gene ID LOC:101499699 (NCBI) by creating a deletion in the coding sequence. The genomic reference sequence is 7071 bp in size with 13 exons (Figure 3A). Desi chickpea cultivar ICC4958 was used in this study. Considering all the parameters, we have selected the 20-nucleotide sgRNA target sequence of *CaPDS* first exon (TGGAGGCAAGAGACGTTCT**AGG**) with the best selection criteria (Supplementary Figure 4). To drive the sgRNA cassettes, we have amplified the promoters of small nuclear RNA *MtU6* of the model legume crop *Medicago truncatula.* A total of eight promoter sequences were amplified from *Medicago truncatula* R108 genomic DNA and cloned in a pGEMT easy vector for further use in chickpea genome editing experiments (Supplementary Text 1). The target sites were selected and cloned under *Medicago* MtU6.1 promoter (508 bp) along with 76 bp Cas9 specific gRNA scaffold and seven bp MtU6.1 terminator flanking with 44 bp adaptor including suitable restriction sites by overlapping PCR as this promoter and its ortholog has been mostly used to edit *Medicago*, and *Arabidopsis* gene (Figure 3B) (Meng *et al.*, 2017; Xing *et al.*, 2014).

**Figure 3:**
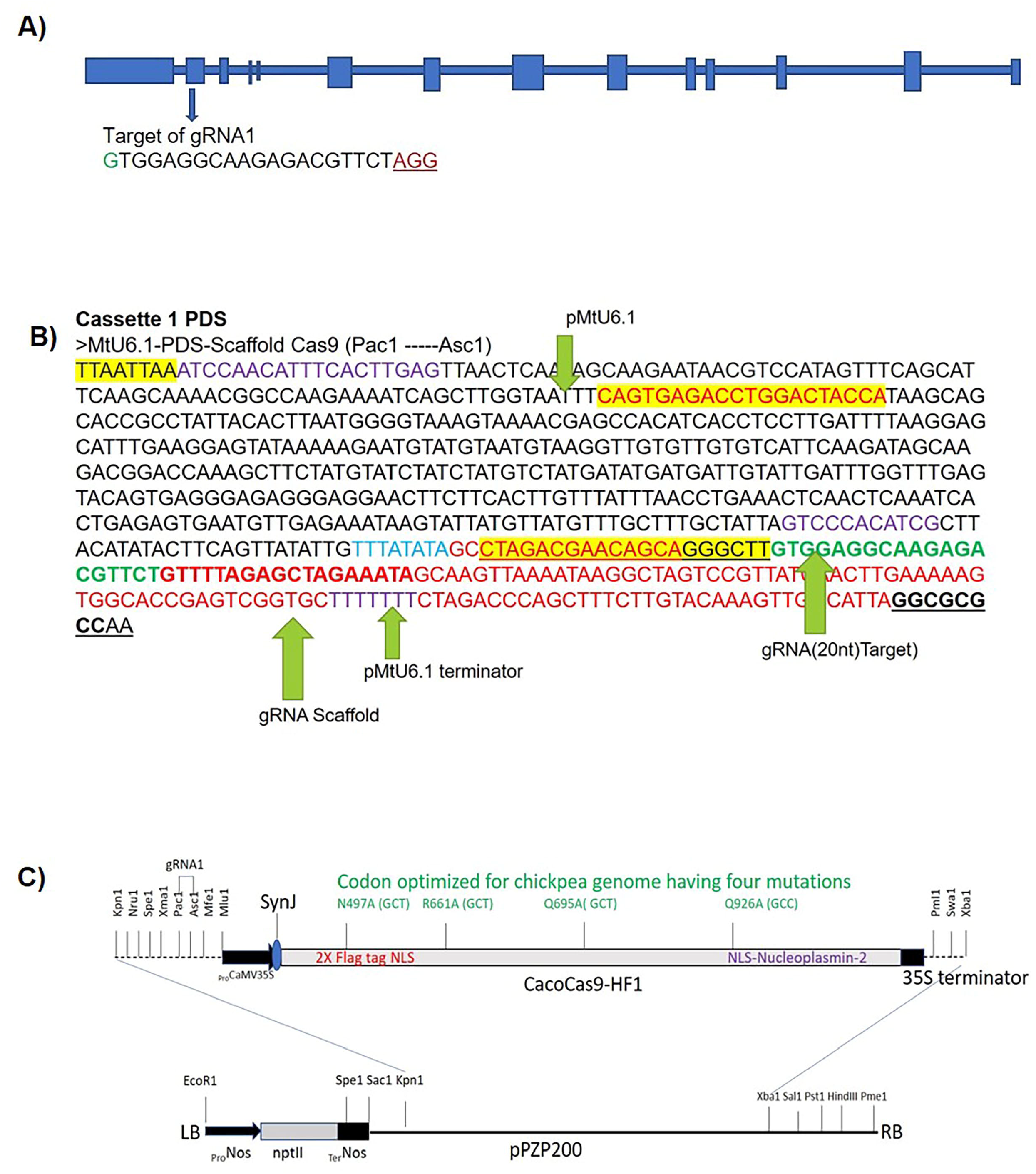
Schematic representations of the chickpea *CaPDS* target gene (LOC101499699). Location of gRNA1 and the CRISPR/Cas9 vector. **(A)** Line representation of chickpea *CaPDS* gene containing 14 exons with the gRNA1 in the second exon. **(B)** Representation of gRNA cassettes for target editing of *CaPDS* gene. The promoter of the small nuclear RNA gene (MtU6.1) was used to drive the expression of gRNA with flanking restriction sites (pac1-Asc 1). **(C)**Schematic map of CRISPR/Cas9 binary vector used for stable *Agrobacterium-mediated* chickpea transformation. The Cauliflower mosaic virus promoter (*CaMV35S)* drives the expression of the Cas9 gene. A total of four mutations were introduced; Arginine to Alaline R497A (GCT), Arginine to Alanine R661A (GCT), Glutamine to Alanine Q695A (GCT) & Glutamine to Alanine Q926A(GCC).

### 3.5 Crop codon optimization of CRISPR/Cas9 and construction of a chickpea transformation binary vector

The modified binary vector *pPZP200* was used in this study by inserting the *neomycin phosphotransferase II;* kanamycin-resistant gene (*NPTII;* EC 2.7. 1.95) as a plant selectable marker, flanking with two restriction enzymes, *EcoR1*, and *Sac1*, respectively. The *nopaline synthase* (nos; EC 1.5.1.19) promoter and the terminator were used to drive and terminate the expression of the *NPTII* gene, respectively. The *pZP200* binary vector containing the *NPTII* gene was further used as a base vector for cloning custom synthesized chickpea codon-optimized Caco*SpCas9*-HF1. *SpCas9* gene was cloned under *CaMV35S* constitutive promoter along with 28 nucleotides synthetic 5UTR (SynJ) enhancer and *35S* terminator sequences. In addition, the *MtU6.1* promoter fused with gRNA scaffold was cloned in the multiple cloning site of the modified *pPZP200* at the PacI-AscI site.

SpCas9 cDNA has been modified and optimized for chickpea codon bias by a tool Gene optimizer (Thermo Fisher Scientific, Inc.) to improve editing efficiency in chickpea. Cas9_CDS_HF1 was optimized and custom synthesized by Invitrogen (Invitrogen Bioservices India Pvt Ltd.) for fine-tuning its expression in chickpea. The oligonucleotide’s GC content was adjusted to prolong the mRNA’s half-life. Codon usage was adapted with a codon adaptation index (CAI) value of 0.72. A total of four non-synonymous mutations (Arg^497^Ala, Arg^661^Ala, Glu^695^Ala, and Glu^926^Ala) were introduced in *SpCas9* to modify it to *CaCoSpCas9* for the enhanced expression in chickpeas (Figure 3C).

### 3.6 *Agrobacterium-delivery* of CRISPR/Cas9 system and phenotypic evaluation of CaPDS edited plants

CRISPR construct was introduced into chickpea single cotyledon half embryo explants through *Agrobacterium-mediated* gene transfer using our modified transformation method. A total of 110 plants were obtained after selection and regeneration. Of these regenerated plants, 48 plants showed albino phenotypes indicating around 43% phenotypic mutant efficiency of the process. Plants grew normally in the tissue culture. According to their phenotypes, these plants were divided into three sets; albino, chimeric, and green. There were 48 albino, 29 green, and 33 mixed phenotypes among the 110 T0 plants. The albino plants were very sensitive to light, and leaves were inclined to die under standard chickpea tissue culture conditions. Therefore, low light intensity was required to maintain these plants in tissue culture for a few days. Even after that, albino chickpea plants did not survive more than 35-40 days. A level of dwarfism was also detected in the albino plants. On subculturing, the albino plants were regenerated as complete albino shoots; the green plants regenerated green shoots, and the mixed phenotype plants showed secondary albino shoots (Table 3; Figure 4A & B).

**Figure 4:**
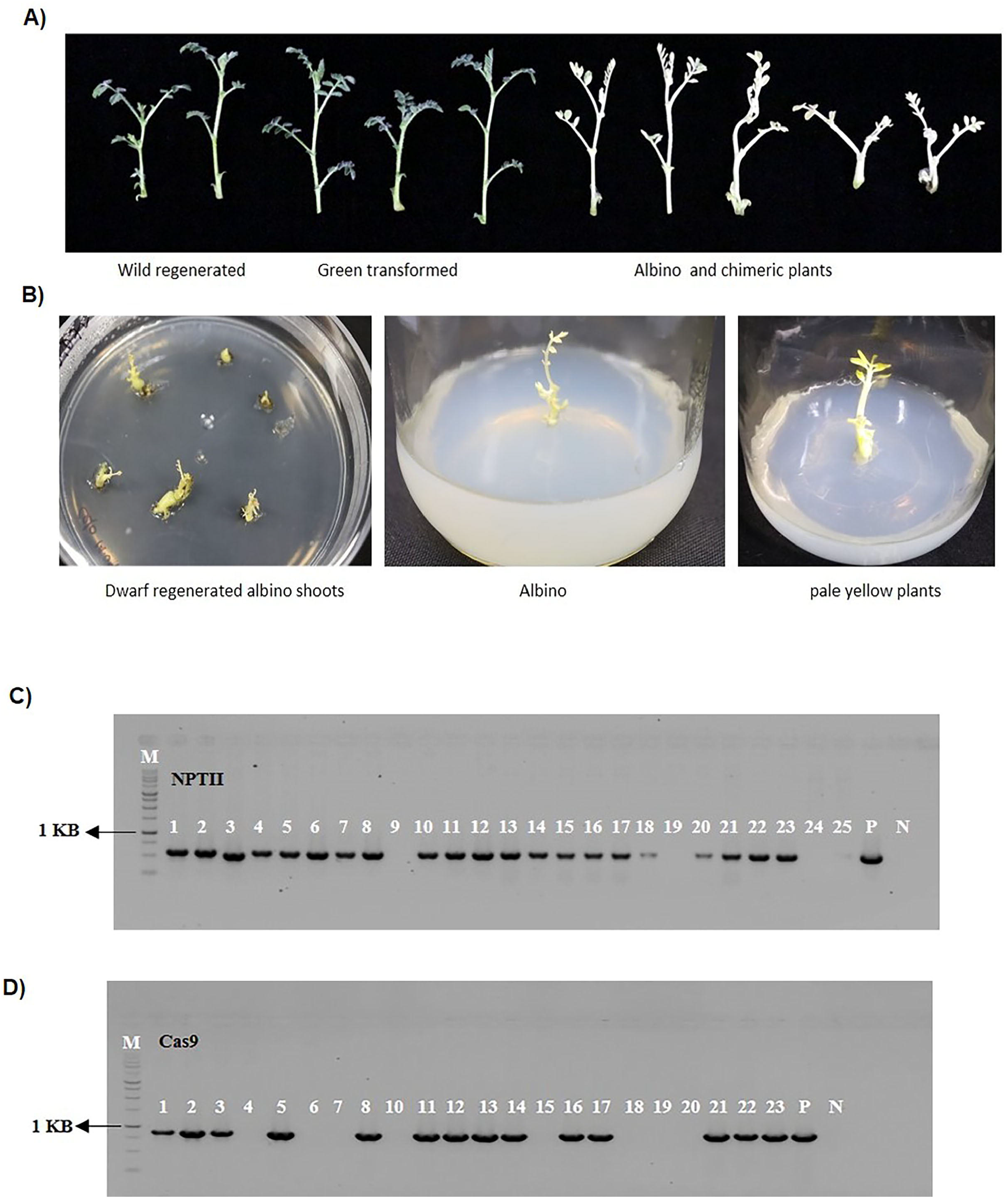
Phenotypes of CRISPR/Cas9-CaPDS_101499699_-mediated mutants of the chickpea. A) Multiple shoots were regenerated from transformed explants. Panels showed albino, chimeric albino, or green with no visible differences compared with non-transgenic wild-type plants. B) Regeneration of albino shoots in tissue culture and their characteristics. C) & D) PCR confirmation of transformants using neomycin phosphotransferase II (kanamycin-resistant gene) and Cas9 gene-specific primers. Genomic DNA was isolated from wild-type regenerated shoots and transformed into regenerated shoots. The sizes of the PCR products are 592 bp (C) and 881 bp (D). Modified vector pPZP200 was used as a positive control (lane P). Lane L, 1 kb DNA ladder. Lane N, negative control.

**Table 3:** Summary of CRISPR/Cas9-mediated mutation types in PDS gene of chickpea plants.

| **Phenotype** | **Albino** | **Chimeric** | **Green** | **Total** |
| --- | --- | --- | --- | --- |
| Number of lines | 48 | 33 | 29 | 110 |
| Percentage of Plants | 43.64% | 30.0% | 26.36% | 100 |
| Number of Sequenced Lines | 21 | 8 | 5 | 34 |

The phenotype percentage was calculated as the number of transformants regenerated each phenotype per 110 lines of T0 chickpea plants.

### 3.7 Targeted mutagenesis and detection by sanger sequencing

To check for the presence of the transgene and investigate any variation in the target sequences, endpoint PCR (Polymerase Chain Reaction) was executed after genomic DNA isolation from 45 regenerated albino plants. Plants were screened with *NPTII* and *Cas9-specific* primers. All the albino plants contain both the *Cas9* and *NPTII* gene integration. Gene-specific primers for *Cas9* (881bp amplicon) and *NPTII* gene (592bp amplicon) successfully amplified the DNA fragments in the transformed chickpea plants (Figure 4C & D).

A 231 bp fragment of the *CaPDS* gene corresponding to the first exon was amplified and sequenced, and mutation analysis was performed with 12 plants showing albino phenotypes. The results obtained from sequencing were typically recorded using nucleotide occurrence chromatograms for particular albino plants. Due to the recalcitrant nature of regeneration and hardening in single-cell culture-based transformation methods, we have developed a protocol for multicellular transformation based on single cotyledon half embryo explants, which results in chimeric plants made up of transformed and non-transformed cells in T_0_ generation. Therefore, Inference of CRISPR Edits of Synthego (ICE) (Conant *et al.*, 2022), an analysis tool for CRISPR, was used to detect mutation in albino plants. The editing efficiency was presented as the indel percentage in the albino plants normalized to that in the untransformed plants. The *CaPDS* sgRNA1 result showed 78%, 69%, 67%, 49%, 39%, 29%, and 19% indel percentage for the albino chickpea plants. We have included only the highest indel percentage (78%) analysis in pictorial representation. The knockout score was 78, which characterizes the proportion of cells that have the mutation. The knockout score was used to know the number of contributing indels expected to result in the functional knockout of the target gene (Figure 5A, B, C & D).

**Figure 5:**
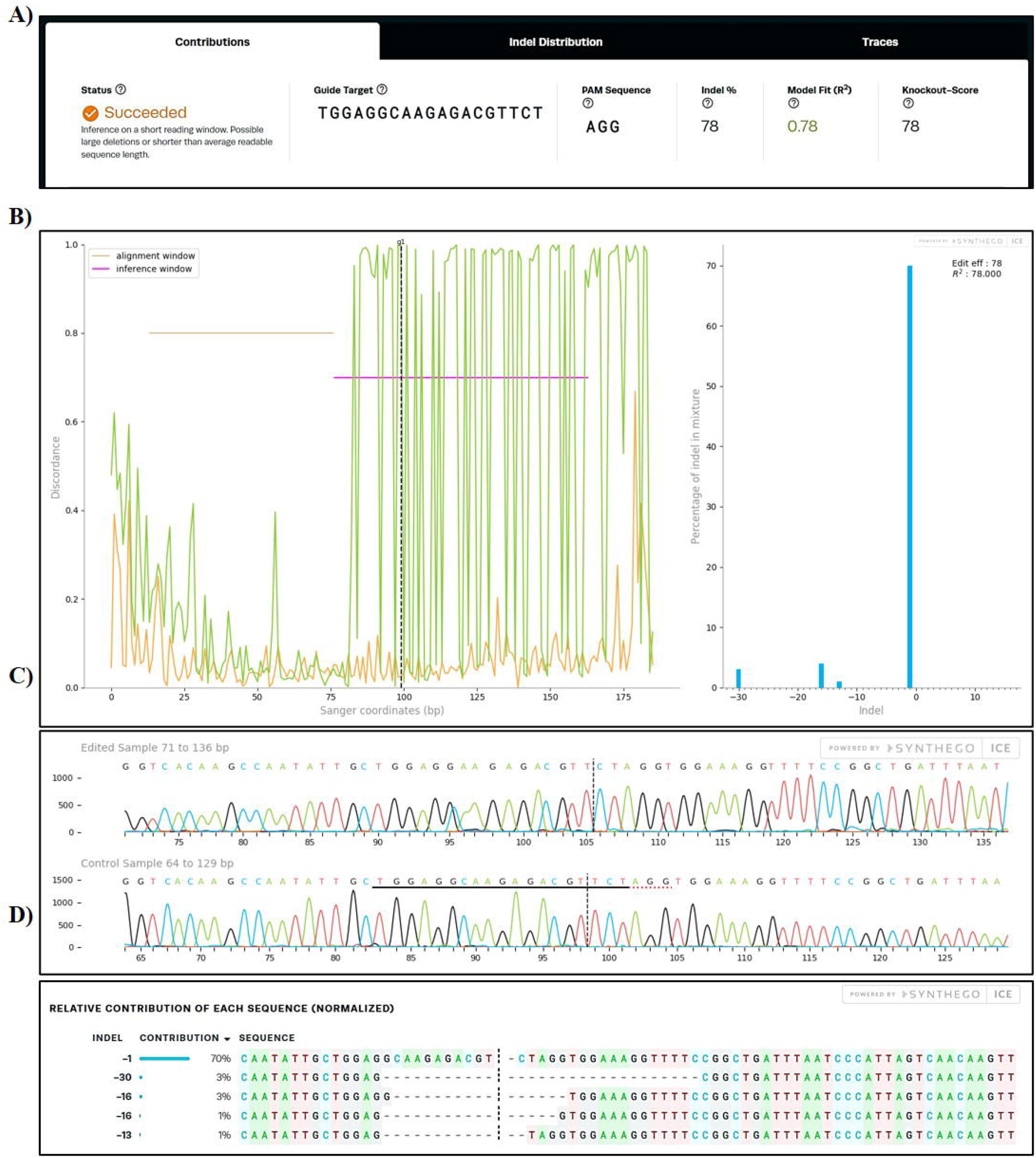
Illustration of the outputs from the ICE (Inference of CRISPR Edits) software for a guide (sgRNA; TGGAGGCAAGAGACGTTCT) targeting the chickpea PDS gene. Transformed chickpea samples were sequenced with Sanger sequencing and further analyzed with ICE. **A)** Representation of ICE outputs generated from sequence files (.ab1) received from Sanger sequencing edited and WT samples. The software output is denoted by a 20 nucleotides gRNA sequence and PAM sequences. Knockout score and InDel percentage received as output post ICE analysis are also exhibited. **B)** Discordance for the edited (green) and control (orange) trace files. The vertical dotted line marks the cut site. The alignment window marks the region of traces with high Phred scores used to align the edited and control traces. The inference window marks the region of the traces around the cut site, which will be used to infer the change in sequence between the edited and control traces. Visualizing this way, we can see that discordance is a robust signal of trace irregularity that can approximate the location and prevalence of editing. As calculated by ICE, insert, or deletion (indel) sizes and their relative prevalence for chickpea PDS gene. **C)** Trace file segments straddling the cut site from the control, and the edited samples are generated for every analysis. The guide target sequence provided by the user is underlined in black, and PAM (Protospacer Adjacent Motif) sequence is denoted by a dotted red underline in the control sample. Vertical dotted lines represent the expected cut sites. **D)** The respective DNA samples of edited and WT chickpea plants were subjected to Sanger sequencing and analysis with ICE. The relative contributions show the sequences inferred in the edited samples and their relative proportions. Vertical dotted lines denote the targeted cut sites.

## 4. Discussion

For more than a decade, *Agrobacterium-mediated* transformation has been employed effectively in grain legumes. (Christou, 1997). There were various studies regarding the generation of transgenic chickpea plants by exploiting *Agrobacterium tumefaciens* (Fontana et al., 1993, Kar et al., 1996; Jayanand et al., 2003; Senthil et al., 2004; Chakaraborti et al., 2006; Hajyzadeh et al., 2015). Earlier, reports were found using *Agrobacterium* strain GV3101 mediated chickpea transformation (Ali et al., 2009). It was reported that cotyledons connected to embryos were essential for healthy shoot generation. Crop improvement in legumes like chickpea through genome editing will require an efficient genetic transformation system that can help to generate numerous transgenic events. Single cotyledon half embryo was used as the explants in these studies, and the plants were established by rooting or grafting. These explants have terminal or axillary meristems that are better sources of totipotent cells (Swathi Anuradha et al., 2008). We have referred to various parameters mentioned in published reports and altered them to an extent that could improve the transformation efficiency through these meristematic embryo explants.

Moreover, dissecting longitudinally one cotyledon attached to half an embryonic axis provides more wounding zones for effective tissue infection with *Agrobacterium* (Polisetty et al., 1997; Chakraborti et al., 2006). It is evident from previous reports on chickpea transformation that earlier methods were largely non-repeatable across most of the laboratories. Few were repeatable with only 0.66% transformation efficiency in chickpea cultivars and further improved up to 2.3% (Bhowmik et al., 2019). In this study, it was observed that on modifying the key steps like exponential increase in selection pressure, devoid of growth regulators during regeneration events and careful hardening of regenerated and selected plants, it was possible to increase the transformation efficiency to about 7.6-fold. Hence, the overall increase in transformation efficiency is due to improvements in key steps and parameters of the earlier published protocol. This modified protocol can be easily used for genome editing experiments in chickpea varieties.

We have used an exponential increase in selection pressure in the current report. The selection system continuously used a step-wise increase in the concentration of selection antibiotic (Kanamycin) at each stage for stringent selection of putative chickpea transformants. The protocol described in this report results in more efficiency and a greater recovery of transgenic plants to produce seeds in a brief time period of 100-120 days.

The CRISPR/Cas9 system has been used for trait development and crop improvement in various crops, including legumes. This study reports CRISPR/Cas9-based codon-optimized gene editing in chickpea plants. Based on published reports on plants, we have made changes for customized synthesis and use the chickpea codon-optimized Cas9. Codon optimization is considered a critical factor in the expression and cellular functions of heterologous genes. Cas9 is derived from bacteria (Streptococcus pyogenes), so to improve its expression and function in plants, the Cas9 gene for plant genome editing has been optimized with plant usage bias codon previously (Jiang et al., 2013; Li et al., 2013; Miao et al., 2013; Shan et al., 2013; Fauser et al., 2014; Xing et al., 2014; Svitashev et al., 2015; Gao et al., 2015; Ma et al., 2015). This study targeted the chickpea PDS gene in transformed stable chickpea plants. The PDS gene is generally used to optimize plants’ CRISPR/Cas9 system (Bae et al., 2021; Hooghvorst et al., 2019; Meng et al., 2017; Fan et al., 2015; Shan et al., 2013). This gene encodes a crucial enzyme in the carotenoid pathway, involving approximately twenty metabolic pathways. These pathways are imperative for the biosynthesis of carotenoid, chlorophyll, and gibberellic acid (Karkute et al., 2017; Pennisi, 2013). Furthermore, the mutation of the PDS gene leads to the generation of photobleached or albino plants, which enables us to identify the CRISPR/Cas9 transformed chickpea plants phenotypically. To avoid confusion and check the efficiency of the codon-optimized Cas9, we have designed and used single gRNA and their delivery in explants through a highly optimized *Agrobacterium-mediated* chickpea transformation method. Chimeric plants showed some green, and mostly the albino phenotypes in the same plants; and this was reported in many earlier studies (Gao et al., 2017; Odipio et al., 2017; Charrier et al., 2019; Pan et al., 2016; Li et al., 2013; Hooghvorst et al., 2019; Bae et al., 2021). Using a single sgRNA in the place of multiple sgRNAs was found better for gene editing and minimized the risk of off-target editing (Yu et al., 2018). Most of the mutations analyzed in edited chickpea plants in this study were deletions.

## 5. Conclusion

Targeted gene editing using CRISPR/Cas9 has facilitated the replacement of transgenic crops possessing exogenous introduced DNA and at the same time allowed to break linkage drag (LD) that limits the trait improvement by conventional or marker-assisted breeding. Site directed nuclease 1 (SDN1), SDN2 and SDN3 can be used alter allele and tissue-specific expression of a trait-linked gene for crop improvement. Therefore, construction of suitable reagents such as, plasmid vectors and optimization of gene editing techniques including the genetic transformation protocol in the popular crops like chickpea would encourage taking-up this crop for agronomic trait improvement through genome editing. We have reported construction of a plasmid vector for gene editing in chickpea by cloning and optimizing different elements within the vector to enhance the efficiency and reproducibility of the technique. We hope that this report will facilitate the use of targeted gene editing in a valuable legume crop, chickpea.

## Author Contributions

**DC:** Initiated, designed, conceived, and coordinated the project. **SKG:** Assist DC in designing, coordinating experimental activities, and writing the manuscript. **NKV:** Molecular cloning, codon-optimized construct development, and molecular screening of transformants. **PM:** Cloning of *MtU6* promoters, gRNA cassettes into a binary vector, and molecular validation of transformants. **PV:** Chickpea transformation and raised edited plants. **NKS:** Imaging of edited plants and performing the supporting experiments. All authors have read and approved the final manuscript.

## Acknowledgements

The authors are thankful to the Department of Biotechnology (DBT), Ministry of Science and Technology, Government of India. DC acknowledges the J.C Bose fellowship (JCB/2020/000014) from the Science and Engineering Board (SERB) – Department of Science & Technology (DST), Government of India. PM acknowledges fellowships from the Council of Scientific and Industrial Research (CSIR), Government of India. In addition, DC is grateful to the DBT-eLibrary Consortium (DeLCON) for providing access to e-Resources. However, the funders had no role in study design, data collection, and analysis, publication decision, or manuscript preparation. We acknowledge Mr. Krishna Kumar and Mr. Bikash Das for their valuable help during various juncture of chickpea transformation, micrografting and hardening.

## Supplementary Information

**Supplementary Figure 1:** PCR amplification of GUS (uidA) and GFP genes from leaves of established plants using specific primers

**Supplementary Figure 2:** Chickpea transformation and selection

**Supplementary Figure 3:** Establishment of regenerated and selected shoots through natural rooting.

**Supplementary Figure 4:** Schematic representation of chickpea *CaPDS* gene with target site indicated as bold, and PAM sequences underlined.

**Supplementary Text 1:** *MtU6* Promoter sequences.

**Supplementary Text 2:** NLS-GFP sequence details.

**Supplementary Table 1:** List of primers used in this study.

**Supplementary Table 2:** *Agrobacterium tumefaciens* (EHA105) mediated the transformation efficiency of chickpea cv. ICC4958

**Supplementary Table 3:** *Agrobacterium tumefaciens* (LBA4404) mediated the transformation efficiency of chickpea cv. ICC4958

## References

Ali, Shahin Sharif, Padma Yealla, and Bidyut Kumar. 2009. “Bar as a Potential Selectable Marker Gene to Obtain Putative Transformants in Indian Chickpea (Cicer Arietinum L.) Cultivars.” Transgenic Plant Journal 3: 2–6.

Badhan, Sapna, Andrew S. Ball, and Nitin Mantri. 2021. “First Report of CRISPR/Cas9 Mediated DNA-Free Editing of 4CL and RVE7 Genes in Chickpea Protoplasts.” International Journal of Molecular Sciences 22 (1): 1–15. https://doi.org/10.3390/IJMS22010396.

Bae, Eun Kyung, Hyunmo Choi, Ji Won Choi, Hyoshin Lee, Sang Gyu Kim, Jae Heung Ko, and Young Im Choi. 2021. “Efficient Knockout of the Phytoene Desaturase Gene in a Hybrid Poplar (Populus Alba × Populus Glandulosa) Using the CRISPR/Cas9 System with a Single GRNA.” Transgenic Research 30 (6): 837–49. https://doi.org/10.1007/S11248-021-00272-9.

Bhowmik, Sudipta Shekhar Das, Alam Yen Cheng, Hao Long, Grace Zi Hao Tan, Thi My Linh Hoang, Mohammad Reza Karbaschi, Brett Williams, Thomas Joseph V. Higgins, and Sagadevan G. Mundree. 2019. “Robust Genetic Transformation System to Obtain Non-Chimeric Transgenic Chickpea.” Frontiers in Plant Science 10 (April): 524. https://doi.org/10.3389/FPLS.2019.00524/BIBTEX.

Briggs, B.A., S.M. McCulloch, and L.A. Edick. 1988. “MICROPROPAGATION OF AZALEAS USING THIDIAZURON.” Acta Horticulturae, no. 227 (September): 330–33. https://doi.org/10.17660/ACTAHORTIC.1988.227.60.

Chakraborti, Dipankar, Anindya Sarkar, and Sampa Das. 2006. “Efficient and Rapid in Vitro Plant Regeneration System for Indian Cultivars of Chickpea (Cicer Arietinum L.).” Plant Cell, Tissue and Organ Culture 2006 86:1 86 (1): 117–23. https://doi.org/10.1007/S11240-005-9072-0.

Chandrasekaran, Jeyabharathy, Marina Brumin, Dalia Wolf, Diana Leibman, Chen Klap, Mali Pearlsman, Amir Sherman, Tzahi Arazi, and Amit Gal-On. 2016. “Development of Broad Virus Resistance in Non-Transgenic Cucumber Using CRISPR/Cas9 Technology.” Molecular Plant Pathology 17 (7): 1140–53. https://doi.org/10.1111/MPP.12375.

Charrier, Aurélie, Emilie Vergne, Nicolas Dousset, Andréa Richer, Aurélien Petiteau, and Elisabeth Chevreau. 2019. “Efficient Targeted Mutagenesis in Apple and First Time Edition of Pear Using the CRISPR-Cas9 System.” Frontiers in Plant Science 10 (February): 40. https://doi.org/10.3389/FPLS.2019.00040/BIBTEX.

Chibbar, Ravindra N., Priyatharini Ambigaipalan, and Ratnajothi Hoover. 2010. “REVIEW: Molecular Diversity in Pulse Seed Starch and Complex Carbohydrates and Its Role in Human Nutrition and Health.” Cereal Chemistry 87 (4): 342–52. https://doi.org/10.1094/CCHEM-87-4-0342.

Christou, Paul. 1997. “Rice Transformation: Bombardment.” Plant Molecular Biology 1997 35:1 35 (1): 197–203. https://doi.org/10.1023/A:1005791230345.

Conant, D., Hsiau, T., Rossi, N., Oki, J., Maures, T., Waite, K., Yang, J., Joshi, S., Kelso, R., Holden, K., Enzmann, B. L., & Stoner, R. 2022. “Inference of CRISPR Edits from Sanger Trace Data.” The CRISPR journal, 5(1): 123–130. https://doi.org/10.1089/crispr.2021.0113.

Fan, Di, Tingting Liu, Chaofeng Li, Bo Jiao, Shuang Li, Yishu Hou, and Keming Luo. 2015. “Efficient CRISPR/Cas9-Mediated Targeted Mutagenesis in Populus in the First Generation.” Scientific Reports 2015 5:1 5 (1): 1–7. https://doi.org/10.1038/srep12217.

Fontana, Giovanna S., Luigi Santini, Sofia Caretto, Giovanna Frugis, and Domenico Mariotti. 1993. “Genetic Transformation in the Grain Legume Cicer Arietinum L. (Chickpea).” Plant Cell Reports 12 (4): 194–98. https://doi.org/10.1007/BF00237052.

Fauser, F., Schiml, S., & Puchta, H. 2014. “Both CRISPR/Cas-based nucleases and nickases can be used efficiently for genome engineering in *Arabidopsis thaliana*.” The Plant journal: for cell and molecular biology, 79 (2): 348–359. https://doi.org/10.1111/tpj.12554

Gamborg, O. L., R. A. Miller, and K. Ojima. 1968. “Nutrient Requirements of Suspension Cultures of Soybean Root Cells.” Experimental Cell Research 50 (1): 151–58. https://doi.org/10.1016/0014-4827(68)90403-5.

Gao, Wei, Lu Long, Xinquan Tian, Fuchun Xu, Ji Liu, Prashant K. Singh, Jose R. Botella, and Chunpeng Song. 2017. “Genome Editing in Cotton with the CRISPR/Cas9 System.” Frontiers in Plant Science 8 (August): 1364. https://doi.org/10.3389/FPLS.2017.01364/BIBTEX.

Gao, Junping, Genhong Wang, Sanyuan Ma, Xiaodong Xie, Xiangwei Wu, Xingtan Zhang, Yuqian Wu, Ping Zhao, and Qingyou Xia. 2015. “CRISPR/Cas9-Mediated Targeted Mutagenesis in Nicotiana Tabacum.” Plant Molecular Biology 87 (1–2): 99–110. https://doi.org/10.1007/s11103-014-0263-0.

Hajyzadeh, Mortaza, Mine Turktas, Khalid Mahmood Khawar, and Turgay Unver. 2015. “MiR408 Overexpression Causes Increased Drought Tolerance in Chickpea.” Gene 555 (2): 186–93. https://doi.org/10.1016/J.GENE.2014.11.002.

Henny, R.J., and W.C. Fooshee. 1990. “Thidiazuron Stimulates Basal Bud and Shoot Formation in Alocasia × Chantrieri André.” HortScience 25 (1): 124–124. https://doi.org/10.21273/HORTSCI.25.1.124.

Hooghvorst, Isidre, Camilo López-Cristoffanini, and Salvador Nogués. 2019. “Efficient Knockout of Phytoene Desaturase Gene Using CRISPR/Cas9 in Melon.” Scientific Reports 9 (1). https://doi.org/10.1038/S41598-019-53710-4.

Jayanand, B., G. Sudarsanam, and Kiran K. Sharma. 2003. “An Efficient Protocol for the Regeneration of Whole Plants of Chickpea (Cicer Arietinum L.) by Using Axillary Meristem Explants Derived from in Vitro-Germinated Seedlings.” In Vitro Cellular & Developmental Biology - Plant 2003 39:2 39 (2): 171–79. https://doi.org/10.1079/IVP2002387.

Jiang, Wenzhi, Huanbin Zhou, Honghao Bi, Michael Fromm, Bing Yang, and Donald P. Weeks. 2013. “Demonstration of CRISPR/Cas9/SgRNA-Mediated Targeted Gene Modification in Arabidopsis, Tobacco, Sorghum and Rice.” Nucleic Acids Research 41 (20): 1–12. https://doi.org/10.1093/nar/gkt780.

Kar, Sanchayita, Debabrata Basu, Sampa Das, Neeliyath A. Ramkrishnan, Puspita Mukherjee, Pritilata Nayak, and Soumitra K. Sen. 1997. “Expression of CryIA(c) Gene of Bacillus Thuringiensis in Transgenic Chickpea Plants Inhibits Development of Pod-Borer (Heliothis Armigera) Larvae.” Transgenic Research 1997 6:2 6 (2): 177–85. https://doi.org/10.1023/A:1018433922766.

Kar, Sanchayita, Tony M. Johnson, Pritilata Nayak, and S. K. Sen. 1996. “Efficient Transgenic Plant Regeneration Through *Agrobacterium-Mediated* Transformation of Chickpea (Cicer Arietinum L.).” Plant Cell Reports 16 (1–2): 32–37. https://doi.org/10.1007/BF01275444.

Karkute, Suhas G., Achuit K. Singh, Om P. Gupta, Prabhakar M. Singh, and Bijendra Singh. 2017. “CRISPR/Cas9 Mediated Genome Engineering for Improvement of Horticultural Crops.” Frontiers in Plant Science 8 (September): 1635. https://doi.org/10.3389/FPLS.2017.01635/BIBTEX.

Krishnamurthy, K. V., K. Suhasini, A. P. Sagare, M. Meixner, A. De Kathen, T. Pickardt, and O. Schieder. 2000. “Agrobacterium Mediated Transformation of Chickpea (Cicer Arietinum L.) Embryo Axes.” Plant Cell Reports 2000 19:3 19 (3): 235–40. https://doi.org/10.1007/S002990050005.

Li, Jian Feng, Julie E. Norville, John Aach, Matthew McCormack, Dandan Zhang, Jenifer Bush, George M. Church, and Jen Sheen. 2013. “Multiplex and Homologous Recombination-Mediated Plant Genome Editing via Guide RNA/Cas9.” Nature Biotechnology 31 (8): 688. https://doi.org/10.1038/NBT.2654.

Li, Wei, Fei Teng, Tianda Li, and Qi Zhou. 2013. “Simultaneous Generation and Germline Transmission of Multiple Gene Mutations in Rat Using CRISPR-Cas Systems.” Nature Biotechnology 31 (8): 684–86. https://doi.org/10.1038/nbt.2652.

Liu, Junqi, Samatha Gunapati, Nicole T. Mihelich, Adrian O. Stec, Jean Michel Michno, and Robert M. Stupar. 2019. “Genome Editing in Soybean with CRISPR/Cas9.” Methods in Molecular Biology (Clifton, N.J.) 1917: 217–34. https://doi.org/10.1007/978-1-4939-8991-1_16.

Liu, Yule, Michael Schiff, and Savithramma P. Dinesh-Kumar. 2002. “Virus-Induced Gene Silencing in Tomato.” The Plant Journal 31 (6): 777–86. https://doi.org/10.1046/J.1365-313X.2002.01394.X.

Ma, Xingliang, Qunyu Zhang, Qinlong Zhu, Wei Liu, Yan Chen, Rong Qiu, Bin Wang, et al. 2015. “A Robust CRISPR/Cas9 System for Convenient, High-Efficiency Multiplex Genome Editing in Monocot and Dicot Plants.” Molecular Plant 8 (8): 1274–84. https://doi.org/10.1016/j.molp.2015.04.007.

Meng, Yingying, Yaling Hou, Hui Wang, Ronghuan Ji, Bin Liu, Jiangqi Wen, Lifang Niu, and Hao Lin. 2017. “Targeted Mutagenesis by CRISPR/Cas9 System in the Model Legume Medicago Truncatula.” Plant Cell Reports 36 (2): 371–74. https://doi.org/10.1007/S00299-016-2069-9.

Miao, Jin, Dongshu Guo, Jinzhe Zhang, Qingpei Huang, Genji Qin, Xin Zhang, Jianmin Wan, Hongya Gu, and Li Jia Qu. 2013. “Targeted Mutagenesis in Rice Using CRISPR-Cas System.” Cell Research 23 (10): 1233–36. https://doi.org/10.1038/cr.2013.123.

Montague, Tessa G., José M. Cruz, James A. Gagnon, George M. Church, and Eivind Valen. 2014. “CHOPCHOP: A CRISPR/Cas9 and TALEN Web Tool for Genome Editing.” Nucleic Acids Research 42 (Web Server issue). https://doi.org/10.1093/NAR/GKU410.

Murashige, Toshio, and Folke Skoog. 1962. “A Revised Medium for Rapid Growth and Bio Assays with Tobacco Tissue Cultures.” Physiologia Plantarum 15 (3): 473–97. https://doi.org/10.1111/J.1399-3054.1962.TB08052.X.

Odipio, John, Titus Alicai, Ivan Ingelbrecht, Dmitri A. Nusinow, Rebecca Bart, and Nigel J. Taylor. 2017. “Efficient CRISPR/Cas9 Genome Editing of Phytoene Desaturase in Cassava.” Frontiers in Plant Science 8 (October): 1780. https://doi.org/10.3389/FPLS.2017.01780/BIBTEX.

Pan, Changtian, Lei Ye, Li Qin, Xue Liu, Yanjun He, Jie Wang, Lifei Chen, and Gang Lu. 2016. “CRISPR/Cas9-Mediated Efficient and Heritable Targeted Mutagenesis in Tomato Plants in the First and Later Generations.” Scientific Reports 2016 6:1 6 (1): 1–9. https://doi.org/10.1038/srep24765.

Pennisi, Elizabeth. 2013. “The CRISPR Craze.” Science. American Association for the Advancement of Science. https://doi.org/10.1126/science.341.6148.833.

Polisetty, R., V. Paul, J. J. Deveshwar, S. Khetarpal, K. Suresh, and R. Chandra. 1997. “Multiple Shoot Induction by Benzyladenine and Complete Plant Regeneration from Seed Explants of Chickpea (Cicer Arietinum L.).” Plant Cell Reports 1997 16:8 16 (8): 565–71. https://doi.org/10.1007/BF01142325.

Popelka, J. Carlos, Nancy Terryn, and T. J.V. Higgins. 2004. “Gene Technology for Grain Legumes: Can It Contribute to the Food Challenge in Developing Countries?” Plant Science 167 (2): 195–206. https://doi.org/10.1016/J.PLANTSCI.2004.03.027.

Qin, Genji, Hongya Gu, Ligeng Ma, Yiben Peng, Xing Wang Deng, Zhangliang Chen, and Li Jia Qu. 2007. “Disruption of Phytoene Desaturase Gene Results in Albino and Dwarf Phenotypes in Arabidopsis by Impairing Chlorophyll, Carotenoid, and Gibberellin Biosynthesis.” Cell Research 2007 17:5 17 (5): 471–82. https://doi.org/10.1038/cr.2007.40.

Saghai-Maroof, M. A., Soliman, K. M., Jorgensen, R. A., & Allard, R. W. (1984). Ribosomal DNA spacer-length polymorphisms in barley: mendelian inheritance, chromosomal location, and population dynamics. Proceedings of the National Academy of Sciences of the United States of America, 81(24), 8014–8018. https://doi.org/10.1073/pnas.81.24.8014.

Sanyal, Indraneel, Aditya K. Singh, Meetu Kaushik, and Devindra V. Amla. 2005. “*Agrobacterium*-Mediated Transformation of Chickpea *(Cicer Arietinum* L.) with *Bacillus Thuringiensis* Cry1Ac Gene for Resistance against Pod Borer Insect Helicoverpa Armigera.” Plant Science 168 (4): 1135–46. https://doi.org/10.1016/J.PLANTSCI.2004.12.015.

Sarmah, Bidyut K., Andrew Moore, Walter Tate, Lisa Molvig, Roger L. Morton, David P. Rees, Pasquale Chiaiese, Maarten J. Chrispeels, Linda M. Tabe, and T. J.V. Higgins. 2004. “Transgenic Chickpea Seeds Expressing High Levels of a Bean α-Amylase Inhibitor.” Molecular Breeding 2004 14:1 14 (1): 73–82. https://doi.org/10.1023/B:MOLB.0000037996.01494.12.

Senthil, G., B. Williamson, R. D. Dinkins, and G. Ramsay. 2004. “An Efficient Transformation System for Chickpea *(Cicer Arietinum* L.).” Plant Cell Reports 23 (5): 297–303. https://doi.org/10.1007/S00299-004-0854-3/TABLES/4.

Shan, Qiwei, Yanpeng Wang, Jun Li, Yi Zhang, Kunling Chen, Zhen Liang, Kang Zhang, et al. 2013. “Targeted Genome Modification of Crop Plants Using a CRISPR-Cas System.” Nature Biotechnology 2013 31:8 31 (8): 686–88. https://doi.org/10.1038/nbt.2650.

Svitashev, Sergei, Joshua K. Young, Christine Schwartz, Huirong Gao, S. Carl Falco, and A. Mark Cigan. 2015. “Targeted Mutagenesis, Precise Gene Editing, and Site-Specific Gene Insertion in Maize Using Cas9 and Guide RNA.” Plant Physiology 169 (2): 931–45. https://doi.org/10.1104/pp.15.00793.

Swathi Anuradha, T., Jami, S., Beena, M., and Kirti, P. 2008. “Cotyledonary node and embryo axes as explants in legume transformation with special reference to peanut,” in Handbook of New Technologies for Genetic Improvement of Legumes, ed. P. B. Kirti (Boca Raton, FL: CRC Press), 253–271. doi: 10.1201/9781439801352.ch17

Tian, Shouwei, Linjian Jiang, Qiang Gao, Jie Zhang, Mei Zong, Haiying Zhang, Yi Ren, et al. 2017. “Efficient CRISPR/Cas9-Based Gene Knockout in Watermelon.” Plant Cell Reports 36 (3): 399–406. https://doi.org/10.1007/S00299-016-2089-5/FIGURES/3.

Upadhyaya, Hari D., Mahendar Thudi, Naresh Dronavalli, Neha Gujaria, Sube Singh, Shivali Sharma, and Rajeev K. Varshney. 2011. “Genomic Tools and Germplasm Diversity for Chickpea Improvement.” Plant Genetic Resources 9 (1): 45–58. https://doi.org/10.1017/S1479262110000468.

Varshney, Rajeev K., Timothy J. Close, Nagendra K. Singh, David A. Hoisington, and Douglas R. Cook. 2009. “Orphan Legume Crops Enter the Genomics Era!” Current Opinion in Plant Biology 12 (2): 202–10. https://doi.org/10.1016/J.PBI.2008.12.004.

Wang, Longlong, Maria Carmen Rubio, Xian Xin, Baoli Zhang, Qiuling Fan, Qiang Wang, Guogui Ning, Manuel Becana, and Deqiang Duanmu. 2019. “CRISPR/Cas9 Knockout of Leghemoglobin Genes in *Lotus Japonicus* Uncovers Their Synergistic Roles in Symbiotic Nitrogen Fixation.” New Phytologist 224 (2): 818–32. https://doi.org/10.1111/NPH.16077.

Xing HL, Dong L, Wang ZP, Zhang HY, Han CY, Liu B, Wang XC, Chen QJ. 2014. “A CRISPR/Cas9 toolkit for multiplex genome editing in plants.” BMC Plant Biology. 14:327–339. https://doi.org/10.1186/s12870-014-0327-y.

Yu, Zhiming, Qiyuan Chen, Weiwei Chen, Xian Zhang, Fengling Mei, Pengcheng Zhang, Mei Zhao, et al. 2018. “Multigene Editing via CRISPR/Cas9 Guided by a Single-SgRNA Seed in *Arabidopsis*.” Journal of Integrative Plant Biology 60 (5): 376–81. https://doi.org/10.1111/JIPB.12622.

