## Supplementary Materials for "Development of an *Agrobacterium*-delivered codon-optimized CRISPR/Cas9 system for chickpea genome editing"

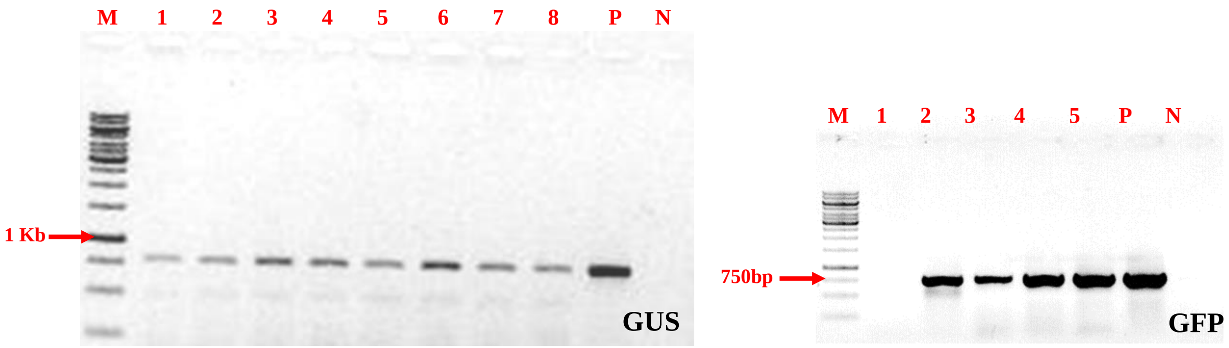


**Supplementary Figure 1:** PCR amplification of GUS (uidA) and GFP genes from leaves of established plants using specific primers (lane 1–6).

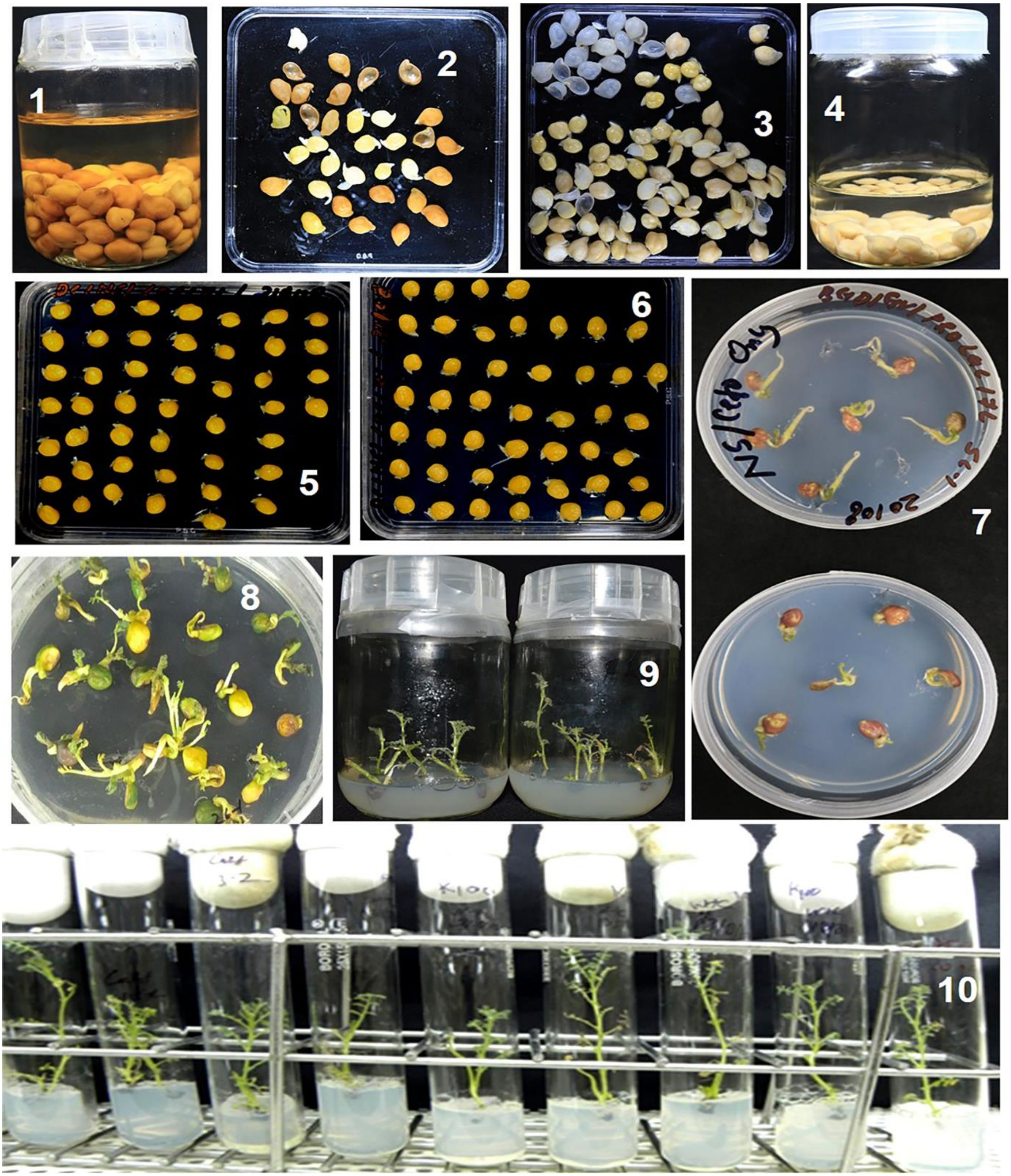


**Supplementary Figure 2: Chickpea transformation and selection.** Preparation of single cotyledon and half embryo explants. These explants were prepared from sterilized and water-soaked desi and Kabuli seeds. Desi chickpea seeds were soaked for 10-12 hours, whereas Kabuli was soaked for 6-8 hours. The seed coats were completely removed. The seeds were intersected along the longitudinal axis with a scalpel to yield two explants (1-4). Each contained half the axis attached to one cotyledon. Shoot regeneration and different selection steps on growth media devoid of growth regulators (6-8). Healthy and elongated shoots were separated from shoot stalk through sub-culturing steps and maintained for further grafting and hardening (9-10).

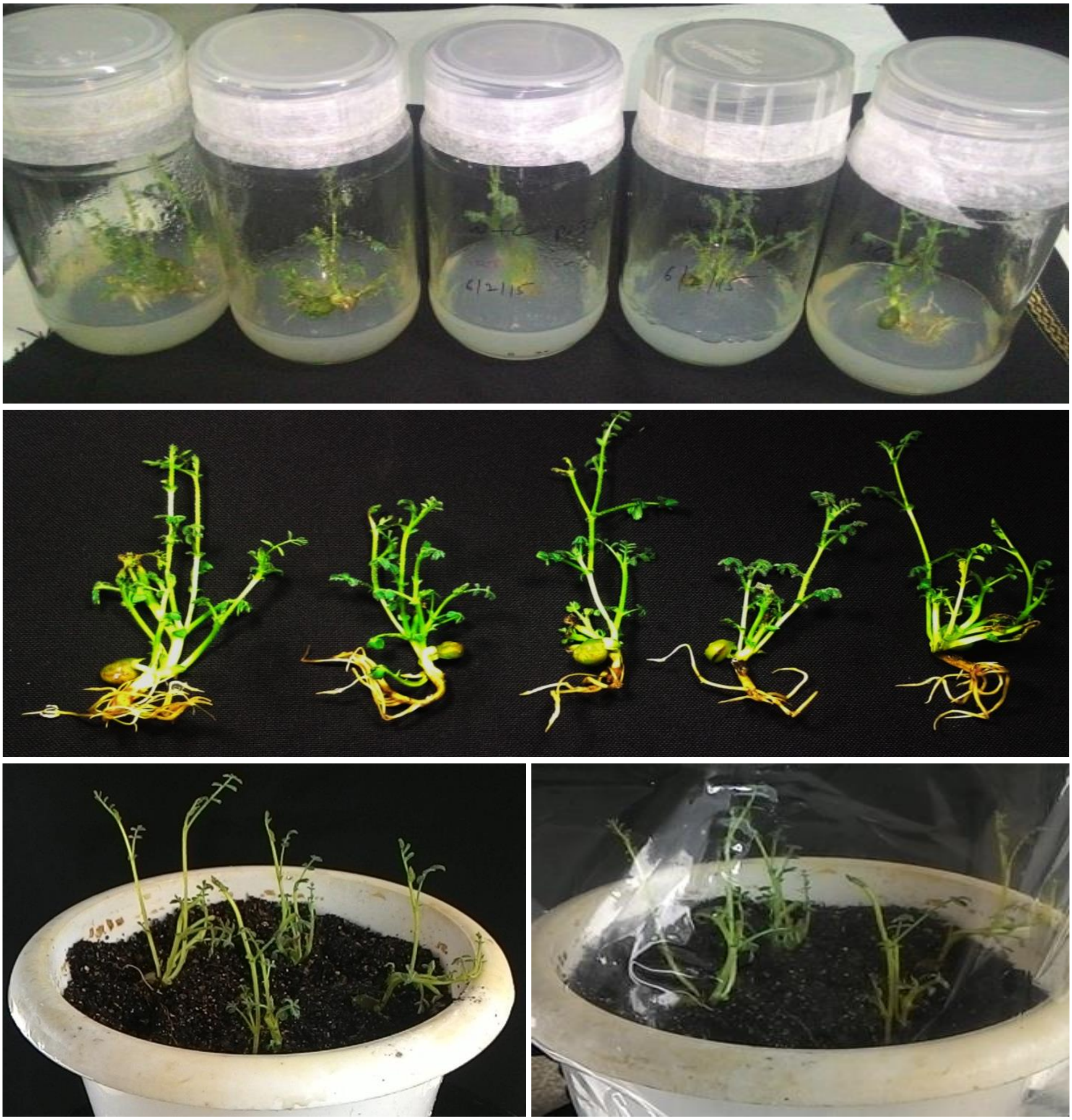

**Supplementary Figure 3: Establishment of regenerated and selected shoots through natural rooting**. **(A)** Development of natural root in marker selected shoots in normal MS media containing selection cefotaxime 250mg $L^{-1}$ devoid of growth regulators and hormones.

aagtcaaaatatttttatataagctataagctattttggatagcttatgaaaattagctcaaaacaacttatagacatatcat
aagctctctcaaatgtgtttatgctagtagataaactcaaataagtaaatcgaaacagacttttttatcacattgtctgagatg
agatccaatgccaataaagaaaaaagttctggaattgtagcattttcaatttttcataaaatttcaaatgaaaggtaccttat
tgtcatgttggttgtctaagcgaaaattgattgtaatatttgcacgagttttggcttctttctcttttggattaatctgtatt
tcaaattatctcagGACTGGCTGGTTTATCAACTGCAAAATATTTGGCAGATGCTGGTCACAAGCCAATATTGCTGGAGGCAA
GAGACGTTCTAGGTGGAAAGgttttccggctgatttaatcccattagtcaacaagtttctgttttatgccttgttttcatcat
ttcagcttgctgttcaattaaatacgaaacttattcttatttgcctgttccttaatataataatggaattgttggttttgtat
gatcaataatacaagttcttttatgaattgaatttctgagactttagagtgggtattgtttaacttcaccatcaagcttgttg
aagattcaacagtgtgtgcaagtttttttattatctcaattttaaatttgttcatactattgtctcgttttctaagGTTGCTGC
ATGGAAAGATGAAGATGGAGACTGGTATGAGACCGGCCTACATATATTCTgtaagttcattaaagtttcaaattagccattgg
cacatgtattgaactattttgatatttctttttagtttTCCAGAAGTCCTTCCATCTCCATTAAATGgtaagatctaatgctt
aatgtatggctgtatattcatataagtataactaccagaagttgaagtctcttaaatgcacatgaaaggccttctatttttaa
aaaagtacaatattggaatttacatttctgctgcaagatgtataaatacgttaataccatctggcaattttcctgcctatgtt
atggctagtgtgc

**Supplementary Figure 4: Schematic representation of chickpea *CaPDS* gene with target site indicated as bold, and PAM sequences underlined.** Exons are shown in capital letters, introns as lowercase, gRNA1 as bold, PAM sequences, and primers used for gene sequencing are underlined.

### Supplementary Text 1

**>MtU6.1 CU013524**

TTAATTAAATCCAACATTTCACTTGAGTTAACTCAATAGCAAGAATAACGTCCATAGTTTCAGCATTCAAGCAAAAC
GGCCAAGAAAATCAGCTTGGTAATTTCAGTGAGACCTGGACTACCATAAGCAGCACCGCCTATTACACTTAATGGG
GTAAAGTAAAACGAGCCACATCACCTCCTTGATTTTAAGGAGCATTTGAAGGAGTATAAAAAGAATGTATGTAATG
TAAGGTTGTGTTGTGTCATTCAAGATAGCAAGACGGACCAAAGCTTCTATGTATCTATCTATGTCTATGATATGATG
ATTGTATTGATTTGGTTTGAGTACAGTGAGGGAGAGGGAGGAACTTCTTCACTTGTTTATTTAACCTGAAACTCAA
CTCAAATCACTGAGAGTGAATGTTGAGAAATAAGTATTATGTTATGTTTGCTTTGCTATTAGTCCCACATCGCTTAC
ATATACTTCAGTTATATTGTTTATATAGCCTAGACGAACAGCAGGGCTTGTCCCTTCGGGGACATCCGATAAAATTG
GAACGATACAGAGAAGATTAGCATGGCCCCTGCGCAAGGATGACACGCACAAATCGAGAAATGGTCCAAATTTTT
TTTGCATTTTTTCCCTTACTCAATCGTTCAGGTTATGTTCTTCCTTTTTTTTTCTGCAAATTAATTTCGTTTTCCACTCG
GTGGTTCCTTTTCTTACTCTTTGATTCTGCGTTTAGGGTTTGTCCAATTTTGACATTGGTGATGT

#### >MtU6.2 AC146705

CTAGAATTCGAACTAAAAAATTTAGACCGTTTGATCAAAGACACGCAGTTCAAATGTGTCTAACAACACCCAAAGT
CAGTGAGAAGTGTGACAAACGGTGAACTTCACCTCGGCTAAGTTTGAATCAAAATAAGAGGACACAAACTTTTAT
GTAGAATATAAGCATATATCAACATAGTGTGACACAAGTAATAGATATTTAAACATTTAGTCGAGAGTTTAATCAC
TGTCAGTGGACATGAAAAAACACTAATTTAGAGTGGTAATAATATCTTAAATAAATCTCAATTAACTCGAGAAAAT
TAGTCTTTACAATTGTGCATGAGAGACCTTTTTATCTAAAAAATTATGAGAATGGAATATGGATATGTACGATTGTA
CTTGTTTCTTGGTCTGAGTGTGAGGGAGAGGAAGAAGTTCACTTAACTTGTTTATTGGTTTATTTAATAAATCTGAA
ACTGAACTCAATTCAGTGAGAGAGCGGGAGATGCATGCATCATGGTCGCTTGCTTTGGACTTGGGTGAAAGCTGG
TTGAGAACTGTATTTGTATATTACGTTTGCTATTAGTCCCACATCGGCTACGTCTTTAGCTTTTAAGCGTTTATATAA
CACACGCGAATATCTAAGCTTGTCCCTTCGGGGACATCCGATAAAATTGGAACGATACAGAGAAGATTAGCATGG
CCCCTGCGCAAGGATGACACGCACAAATCGAGAAATGGTCCAAATTTTTTTTTGACTTTTTTTTTCCTTCAATTTCAC
ACTCGTAAGTTTTCTCCGTTCAGGTATGATTTCTTTTCTGCATTTGGTTTTCAATTCCGTTCCTTTTCACTTAGATTTTT
GTGTTTAGGGTTTTGTTTTCGTTTCTGTAAGTATTGACGAGGTGCTGATGACAATTT

#### >MtU6.3 AC140026

TTAATCACATCAATTGATATATTATAATTTTATTAATTTTATTTTTCTTGGTTTTGTGTATGTGCCTAAAATTGAAGTG
TTGGATATGTGTCCAACCAAGACTTTCTCATGGCCAAACACTCATATTTGAGCACAAATTTCATTACAATGATTAAG
ATTACAATAAAACCCTTTGATTTATTTGGATGAGAAAGTAAATAAAAATCCAATAAATGAAACAGAAAAACAGACA
GTGATATAAATAATATATAAAGATGATAAAAATAAGGTGGCAAGCTAAGAAAAAAAAAATTCGAGCGGTGAAAGG
GATAGTGTCTAGTGGTTCAGACTACACTAACTTGAGGTACCTATACCTAATATTTTGTTTAAAATAAAAAGAAAAG
AATAAAGAAAACAAAAGATCATCGTTTTTGTTGTGTAATGTTACTCTGTAGGAAGAAATTTCTATTTGCTATGGACT
GGAATTTCGAGCAAGAGCTCATTGAGAAAATGCATCTTTCGCAGTAAGAGGTGCGTGTTGTTTGTTTTTGTCCAGA
GTAAGTCCCACATCGACTAAATATGTGAAAATTTCAGTCTTTATATTACGCTACAACACCTCCTTGCTTGTCCCTTTG
GGGACATCTGATAAAATTGGAACGATACAGAGAAGATTAGCATGGCCCCTGCGCAAGGATGACACGCACAAATC
GAGAAATGGTCCAATTTTTTTTTTTTATTGGTTTGTGTTTTTGTGGCTATTTTCATTTTCAATTTCGTGACGTCTCCTTT
ATGTTGCAGATGTAGTACCTTAACATAGTTTGTGTGAGTTTGAATGAATAAGGACGATACATGTTTCCAGATTTTTT
ATTCACACAGGTCTTCATATAATTTGATTCTGTTTTGTCCCCTTGATTTCAGTTTGTAGAGTAGGTTTAGGAGAGGA
ATTAGATGCTAATCTTGTCTTGCATACTCACCTATGAAGCTCTGACACAAACATTGACACATCAACGCCGATGATAA

TTTGATAAAATTAATGAAATGAATGTAGTCACATGTGTCAATGTCATGTCGGAGATACCAACATGCATCGGACACC
GGACAGTTCGATCAAAGTGTCGGTGTTACATAGATACTCACTAATGTATGAACACTATAGTGAGTGT

#### >MtU6.4 AC174349

AGGAAATGAAGAATGGAGGTGGTGGTGGTTCAAATCCGGCCATGGCGTCACCGGATTCAGAGAAATGTTGGCTG
TTTATTTTCAATACATGTGCAAATTGCAAGCGAGCAATTTCCAGATACTTTCTCTAACATCTCTCTTGGTTGAACAAT
CTCCCTAACCTAATTCAAATTTATTTCCTCCACACCAAAACAAAGCATTGAATGATTTAATTTCAGTTTAAGAAAAGT
TTTGATGGAAAGGGGCAATAAAAAATATACATGATTTCATCTTGTCCATCTTTATTCTTTATCAATGGCAGAGAAGA
GAAAACTAACTTGACATAAATTACAAACATAAAATTTAACCATGTTGAATTAAATTACAAAATTGATTTTCCCTTTAA
CACAGTTGAATTCTGTTGTAATTGGGTAGGATTAGAAAAGCTTTGATGGAAAAGTGGTGGTGATTGTGCGGTGGG
GAATTGGTGAAGACTTGCAAGCTGTTTTTTCTCTTTTTATATATATATATATATATATTTTTTTTTTACCTTTTTTCTAG
AGATTAATCTAAAGGCTAGGAATAAGGTGGTACAAAGGTGGTATATGAATAAGGTGTCTACTTATAATTATCCTTA
ATTTATACATCTATCTACATCACCATTATATCAAAACAATATGAAAGACAAGTAGCAAAAACATTTCTCTCTCATATA
TCTCTCGACTCCACATAAAGGATATAATTCTTCACTCTTAATACCTTTCACTCTATTTAAATGATAAAAAATATTGTCT
TTATTTTTTACTCACTTTTCCATGGTTTAATTATCTTTCTGTATATTTGTGAAAAACATTATCTCACTGTATTCAAATTA
GCAAAGGACAACTTTTGAAGATTCCTTTAAAGATATTTGTTTTCATTATCGATATTAAAATATTATAGTACTAGTTGA
CACTTAAAGAACTGAAAACTGGTGGCCTCTAAACCTTTCTGAATGTGTAGCTGGTTTGACAACCATGGAAGGGTAC
GAACTTTTGAAGTTTATTTTATTTTAAAAATGATCCGATTTTTTTTTTAATTTTGGGAAATAACATCAACAAG

#### >MtU6.5 AC133780.33

TTTCAATGTAGAATTATTATTTTACTTCCAATTATACCCTCTAATTAATTTCCAAAGCATTATACCAATAATAAATAAA
GTTAGTTTAGTAAAATTGTCATATCATTTAACATTATTATTAGATTTCTTAATTTGTGTTTAAAAGCTTTAAACGATG
ATCATTTTTAAACAGAGAGTATAAAGTAGTAAAATAGTACTATTAGAAATGAATTGACGTGACATGCTATGAAAAG
TCTGGAAGAGTATCGATAAAAGGCTACACTAGAGGTAGCTACTTATATGCGCAGGAACTGAAATCAAAAATAAAA
TAAAGAAGAAGGAAGATGCATGTTGTGTTATATAAGTGAAGGAGAAGGACTTGCATGTTGTGTTATATTTGCTTG
TTTTAGTCCCACATCGACTGAAACAGAAAATATCTAGGCGTTTATATACTGCAAGCGAACCATAATGTTTGTCCCTT
CGGGGACATCCGATAAAATTGGAACGATACAGAGAAGATTAGCATGGCCCCTGCGCAAGGATGACACGCACAAA
TCGAGAAATGGTCCAAATTTTTTTGATATTTTTTTCTTCAAATTTTCTCTTTTCTGAGCTCAGATAGGTTCTTCTTTCTT
CATTTTGTTTTCCAGATTAGGTTTTTATGACCATTTTGAAATTGGGGGTGTCTCCTTTACTGTTAAACCAGTATCAGC
CTATATTATTTCCCTGTTTATTTTTATTTTTTATGATCCTAAATGTATAGTT

#### >MtU6.6 AC133780

TTGATTTAGCTAAACTTGGGTTATTTGAAGAAGCAGATGAGCTATCAATGAGATCTACATTTGCAGAGTCAGTTTA
CAATGATGTTGTTTTTGGAATATGCCTATCTTATATGATCAATGAGGCATTTAATTGGGTGCATATGATGGTGAAAA
AAGGTGCAGCTCCTGGCTTGGGAATGATGACTCATGTGGAATTTGGTCTTAAATTTATCACATCCTTTTGGGATGT
GATGATTGTATCACTTGTTCATTTTGCAAAGACAAGGTGCACTGCTACAAACTTTGGTTTAATCTGAAATAAAACAA
AACTCACTGAGAGGAAGATGCATCCCAGTAGGTGAAAGTCGAGAAGGATTTGCATGTTACTATTACACTTGCTTTT
TAGTCCCACATCGTCTGAAACAGAAAATATTTCAGCGTTTAAATACTTCAAGCGAACCAGTAGGCTTGTCCCTTCGG
GGACATCCGATAAAATTGGAACGATACAGAGAAGATTAGCATGGCCCCTGTGCAAGGATGACACGCACAAATCG
AGAAATGGTCCAAATTTTTTTGCCATTTTTTTCTTCGAAATTTCTCTTGTCTGAGTTCAGATAGGTTCTTCTTTCAGCA
TTTTGTTTTCCATATTAGGTTATCTTGTCACTTTTGAAATGGGGGGGGGGGGGGGGGTGTTTCTTACTGTTAGATTA
ATATCAGCCTTTATAGTACTGTATTCTTACCAAAGTATTACAACATCTATCCTTGTTTATTTTTATTTTTTGTGATCCT
AAATGTATAGTCTGATTTGGTTTAAGATATATACAAAATTTTAATCT

#### >MtU6.7 AC133780

ATAGTGGAGTTAGCGATTAGTTGCATTTCAGGGAAAAGGGTCAACTTTTTGTTGATTCTATTGGATATTAGTTGAG
AGTTGAGACTATGTTGTAGGCTGGAGATTTTTCAACTCCGTGTTTGTTTCTGGGGGTGGTGCTATTGAAGCTATTG
TAGTAGGATGAATCGATATCTGCTATAGTGATCAATAGGGTAATGAAATCAAATTATATAAATTCACCGACGGCAC
GAAAGTGTGGTTACTTGACAACAGGGAATCGATATCCGTTATAGTGATCAATAGGGTAATGAGCATATTACAGTTT
GCTTTGTATTACAGTTAGTATATGTTGTGACAAACTTTGGTTCAATCTGAAATAAAACTCAACCCACTGAGAGGGA
GGTACATCTGAGTCGCTTGCTTGCTTCAGGTGAAAGTTGAGAAGGATATGCATATTACTATTACACTTGCCTTTTAG
TCCCACATCGGCTGAAACAGAAAATATTTTAGCATTTATATAGAGCAAGCGAACCGTAAGGCTTGTCCCTTCGGGG
ACATCTGATAAAATTGGAACGATACAGAGAAGATTAACATGGCCCCTGCACAAGGATGACACGCACAAATCGAGA
AATGGTCCAAATTTTTTTTGACATTTTTTTCCTTCAGAATTTCTCTTTGATTTTTACTCATT

#### >MtU6.8 AC133780

CAGAAATCATTTCCACTGCAAAATGGTTAATGGGCTGGGATTTATGCAGGGTGCATAAGGATTCCTTAT
GCATAATAACCGCACCCATTAGGAGCCTATGACAGTTGAAGCTGTGCTTGTTGAAGCATTGGGATTGAG
ATGGTGATTCCTAAAAGCAAAGGACTTAAAACTCCAAAGGGTCAGTGTAAAATCTGATGCAGCTGTTGT
AGTAAACTGCATCAATTGCATCGACTTGGAGATGTAATTAGAAGGTAATTCATATGTGAAAGATGAAGA
CTCTTATGGATGCGTAAGATTTTGTTTTAGTGGATATTTAATCTTTTAACTTGCTTAAAAATTAACTGCAG
CACTAAGCATCGAGTAAGACACAAATTGTGAGTGAATCATTATGAAAAATAGGGAAAGGGACACTCCA
GCTTTAGTTTAAATAGAGGATCACACTCTTGTATCCTGTGTGGCAAAGCAACATAAACCCTTTCCCTTAAC
ATACTTTGGCTCTCTTTATAGGAATCGATATCCGTTATAGTGATATCTGAATCGCTTGCTTGCCTCAGGTG
AAAGTTGAGAAGGATTTGCATGTAACTGTTACACTTGCTTTTTAGTCCCACATCGACTGAAATAGAAAAT
ATTTTGGCGTATATATAGAGCAAGCAAACCATAAGGCTTGTCCCTTCGGGGACATCCGATAAAATTGGA
ACGATACAGAGAAGATTAGCATGGCCCCTGCGCAAGGATGACACGCACAAATCGAGAAATGGTCCAAA
TTTTTTTTGACATTTTTTTTCTTAAGAATTTCTCTTTGATTTGGGGATAATTTGGTTCAAGTTAATTATTCTG
AAG

**Supplementary Text 2**

**>NLS-GFP**

ATGCCTAAGAAGAAGCGTAAGGTTGGAATCGTGAGCAAGGGCGAGGAGCTGTTCAC
CGGGGTGGTGCCCATCCTGGTCGAGCTGGACGGCGACGTGAACGGCCACAAGTTCA
GCGTGTCCGGCGAGGGCGAGGGCGATGCCACCTACGGCAAGCTGACCCTGAAGTTC
ATCTGCACCACCGGCAAGCTGCCCGTGCCCTGGCCCACCCTCGTGACCACCTTCACC
TACGGCGTGCAGTGCTTCAGCCGCTACCCCGACCACATGAAGCAGCACGACTTCTTC
AAGTCCGCCATGCCCGAAGGCTACGTCCAGGAGCGCACCATCTTCTTCAAGGACGA
CGGCAACTACAAGACCCGCGCCGAGGTGAAGTTCGAGGGCGACACCCTGGTGAACC
GCATCGAGCTGAAGGGCATCGACTTCAAGGAGGACGGCAACATCCTGGGGCACAAG
CTGGAGTACAACTACAACAGCCACAACGTCTATATCATGGCCGACAAGCAGAAGAA
CGGCATCAAGGTGAACTTCAAGATCCGCCACAACATCGAGGACGGCAGCGTGCAGC
TCGCCGACCACTACCAGCAGAACACCCCCATCGGCGACGGCCCCGTGCTGCTGCCC
GACAACCACTACCTGAGCACCCAGTCCGCCCTGAGCAAAGACCCCAACGAGAAGCG
CGATCACATGGTCCTGCTGGAGTTCGTGACCGCCGCCGGGATCACTCACGGCATGGA
CGAGCTGTACAAGTAA

**Supplementary Table 1**

| Primer | 5’>>>>>>3’ |
|---|---|
| Cas9_F | CTAGTATACTAAACCATGGACTACAAGG |
| Cas9_R | AAGTTGCTCTTGAAGTTTGGGGT |
| pZP200 seq_F | TGGCAGGATATATTGTGGTGTAAAC |
| pZP200 seq_R | GTTTACCCGCCAATATATCCTGTCA |
| NLS-GFP_F | CACCATGCCTAAGAAGAAGCGTAAGGTTGGAATCGTGAGC |
| NLS-GFP_R | TTACTTGTACAGCTCGTCCATGCCGTGAGTG |
| MtU6.1 Set1_F | GGTTAATTAAATCCAACATTTCACTTGAG |
| MtU6.1_PDS _R1 | TCTCTTGCCTCCACAAGCCCTGCTGTTC |
| MtU6.1_PDS _F1 | GTGGAGGCAAGAGACGTTCTGTTTTAGAGCTAGAAATA |
| U6.1ter_UnivRAsc1 | GGCGCGCCTAATGCCAACTTTGTAC |
| SEQ_F_PDS1_Frag1 | ATATTTGCACGAGTTTTGGCTT |
| SEQ_R_PDS1_Frag_1 | TTGAACAGCAAGCTGAAATGAT |
| Npt_II_F | GATCTCCTGTCATCTCACCTTGCT |
| Npt_II_R | GCCTTCTTGACGAGTTCTTCTGA |
| MtU6.2_F_Full | CAACACCCAAAGTCAGTGAG |
| MtU6.2 R_Full | TATCGGATGTCCCCGAAG |
| MtU6.3_F_full | ACCAAGACTTTCTCATGGCC |
| MtU6.3_R_full | GGAGGTGTTGTAGCGTA |
| MtU6.4_F_full | ACCAAGACTTTCTCATGGCC |
| MtU6.4_R_full | GGAGGTGTTGTAGCGTA |
| MtU6.5_F_full | TACTTCCAATTATACCCTCTA |
| MtU6.5_R_full | TATGGTTCGCTTGCAGT |
| MtU6.6_F_full | ATGCCTATCTTATATGATCAATG |
| MtU6.6_R_full | ACTGGTTCGCTTGAAGT |
| MtU6.7_F_full | GGTGCTATTGAAGCTATTGTAGT |
| MtU6.7_R_full | CTTACGGTTCGCTTGCTCT |
| MtU6.8_F_Full | TGAAGCATTGGGATTGAGATGG |
| MtU6.8_R_Full | CTTATGGTTTGCTTGCTCT |
| GFP_F | ATGGTGAGCAAGGGCGAGG |
| GFP_R | TTGTAGTTGCCGTCGTCCTTGAA |
| GUS_F | CTGTGGGCATTCAGTCTGGA |
| GUS_R | TCTGCCAGTTCAGTTCGTTGTT |

#### Supplementary Table 2: *Agrobacterium tumefaciens* (EHA105) mediated the transformation efficiency of chickpea cv. ICC4958

| **Construct used** | **Explant type** | **No. of explant** | **No. of regenerated shoots (After cocultivation)** | **No. of a kanamycin-resistant single shoot on different concentration** | | | | | | **Average Transformation frequency (%)** |
|---|---|---|---|---|---|---|---|---|---|---|
| | | | | **Kan 50** | | **Kan 100** | | **Kan 200** | | |
| | | | | **I** | **II** | **I** | **II** | **I** | **II** | |
| **ProCaMV::GUS** | Half embryo longitudinally with one cotyledon | 1850(15) * | 836 | 698 | 459 | 345 | 271 | 221 | 178 | 9.62 |
| **ProCaMV::GFP** | Half embryo longitudinally with one cotyledon | 1498(12) * | 543 | 477 | 333 | 256 | 183 | 162 | 108 | 7.02 |
| | Total | 3348(27) * | 1397 | 1175 | 792 | 601 | 454 | 383 | 286 | 8.54 |

*No. of independent transformation events

#### Supplementary Table 3: *Agrobacterium tumefaciens* (LBA4404) mediated the transformation efficiency of chickpea cv. ICC4958

| **Construct used** | **Explant type** | **No. of explant** | **No. of regenerated shoots (After cocultivation)** | **No. of a kanamycin-resistant single shoot on different concentration** | | | | | | **Average Transformation frequency (%)** |
|---|---|---|---|---|---|---|---|---|---|---|
| | | | | **Kan 50** | | **Kan 100** | | **Kan 200** | | |
| | | | | **I** | **II** | **I** | **II** | **I** | **II** | |
| **ProCaMV::GUS** | Half embryo longitudinally with one cotyledon | 1850(15) * | 406 | 307 | 240 | 198 | 144 | 118 | 90 | 4.86 |
| **ProCaMV::GFP** | Half embryo longitudinally with one cotyledon | 1498(12) * | 410 | 360 | 258 | 204 | 182 | 137 | 92 | 6.14 |
| | Total | 3348(27) * | 81 | 667 | 498 | 402 | 326 | 255 | 182 | 5.43 |

*No. of independent transformation events
